# Neural classifiers with limited connectivity and recurrent readouts

**DOI:** 10.1101/157289

**Authors:** Lyudmila Kushnir, Stefano Fusi

## Abstract

For many neural network models in which neurons are trained to classify inputs like perceptrons, the number of inputs that can be classified is limited by the connectivity of each neuron, even when the total number of neurons is very large. This poses the problem of how the biological brain can take advantage of its huge number of neurons given that the connectivity is sparse. One solution is to combine multiple perceptrons together, as in committee machines. The number of classifiable random patterns would then grow linearly with the number of perceptrons, even when each perceptron has limited connectivity. However, the problem is moved to the downstream readout neurons, which would need a number of connections that is as large as the number of perceptrons. Here we propose a different approach in which the readout is implemented by connecting multiple perceptrons in a recurrent attractor neural network. We prove analytically that the number of classifiable random patterns can grow unboundedly with the number of perceptrons, even when the connectivity of each perceptron remains finite. Most importantly, both the recurrent connectivity and the connectivity of downstream readouts also remain finite. Our study shows that feed-forward neural classifiers with numerous long range afferent connections can be replaced by recurrent networks with sparse long range connectivity without sacrificing the classification performance. Our strategy could be used to design more general scalable network architectures with limited connectivity, which resemble more closely the brain neural circuits which are dominated by recurrent connectivity.

## 1 Significance statement

The mammalian brain has a huge number of neurons but the connectivity is rather sparse. This observation seems to contrast with the theoretical studies showing that for many neural network models the performance scales with the number of connections per neuron and not with the total number of neurons. To solve this dilemma, we propose a model in which a recurrent network reads out multiple neural classifiers. Its performance scales with the total number of neurons even when each neuron of the network has limited connectivity. Our study reveals an important role of recurrent connections in neural systems like the hippocampus, in which the computational limitations due to sparse long range feed-forward connectivity might be compensated by local recurrent connections.

## 2 Introduction

The performance of a neural circuit is often evaluated by determining the number of input-output functions that can be implemented, or equivalently by the number of inputs that can be classified correctly by the neural circuit. Theoretical studies on perceptrons [1] and recurrent neural circuits (see e.g. [2]) have shown that typically the performance of a neural circuit scales with the number of synaptic connections that individual neurons receive, and not with the total number of synapses, or with the total number of neurons (see e.g. [3]). This is clearly a problem in the biological brain in which the connectivity is sparse, especially when long range connections are considered [4]. One striking example is the mammalian hippocampus [5]. A typical pyramidal neuron in rodent CA3 receives only 50 synapses from the upstream area[6], the dentate gyrus (DG), which contains around 10^6^ neurons. Not only the connectivity is sparse but also the neural activity[7, 8].

One possible way to overcome the limitations of sparse connectivity is to adopt the strategy of “committee machines” [9], which are basically populations of classifiers. Each classifier is weak, as a perceptron with limited connectivity, but the output is generated by reading out a large number of weak classifiers and by combining them together using a majority vote or some more sophisticated strategies. The final classification performance is significantly better than the one of each individual classifier, provided that the errors of the individual classifiers are sufficiently independent. The term committee machines goes back to 1960-s [9], but they have also been a focus of more recent studies (see e.g. [10], [11]) and basically they are all based on strategies that in machine learning are known as *ensemble methods* or *hypothesis boosting* [12, 13], strategies that are often adopted also in statistics [14, 15, 16, 17]. Some of the examples include stacking [18, 19], bagging [20], arcing [21] and adaboost [22, 23].

One class of committee machines is implemented using populations of neurons, each essentially behaving as a neural classifier, like a perceptron [24, 25, 26, 27]. Classifiers with limited connectivity are weak classifiers. It is possible to compute the classification capacity when each neural classifier has sparse connectivity [25]. The connections between the *N* input neurons and the *M* < *N* neural classifiers are assumed to be non-overlapping (*N/M* connections per “perceptron”) and plastic. The final response of the committee machine is obtained by majority vote of the *M* neural classifiers, which can be easily implemented by introducing a readout neuron that is connected to all the neural classifiers with equal weights. The maximum number of correctly classified inputs is proportional to 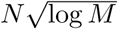 whereas each neural classifier would not go beyond *N/M* inputs. This is a favorable scaling and it is similar to the one obtained in other committee machines. However, one has to keep in mind that in these implementations the neural classifiers have sparse connectivity, but the readout neuron performing the majority vote should have a number of connections that scales with *N*.

Here we propose a network architecture that overcomes the restrictions imposed by the limited connectivity, as in the committee machines, but it replaces the readout neuron that has extensive connectivity with a more biologically plausible recurrent network in which all the neurons have a number of connections that remains finite when the number of classifiable patterns grows unboundedly. More specifically we show that the number of random inputs that can be correctly classified scales linearly with the number of input neurons *N*, even when the number of connections per neural classifier *C_F_* does not increase with *N*. The number of neural classifiers *M* is assumed to be proportional to *N*.

Interestingly, under certain conditions the recurrent scheme has larger classification capacity than the majority vote scheme. This happens for sparse input representations, the regime that is relevant for the mammalian hippocampus and that we investigate in detail.

## 3 Methods

### 3.1 Fully connected readout

In this section we derive the classification capacity of a single fully connected linear threshold readout, or *perceptron* (see Figure 1a) achieved with a simple learning rule that we employ throughout this work. We assume that the input patterns and labels are random and uncorrelated, meaning that the activity of each input unit as well as the label for each pattern is chosen independently, which makes calculations analytically tractable. We use a simple Hebbian-like learning rule, that is not optimal, and thus leads to a lower capacity than Cover’s 2*N* result [28]. However, the scaling of the maximal number of learned input patterns *P*_max_ with the number of input units *N* is still linear, as is shown below.

**Figure 1.**
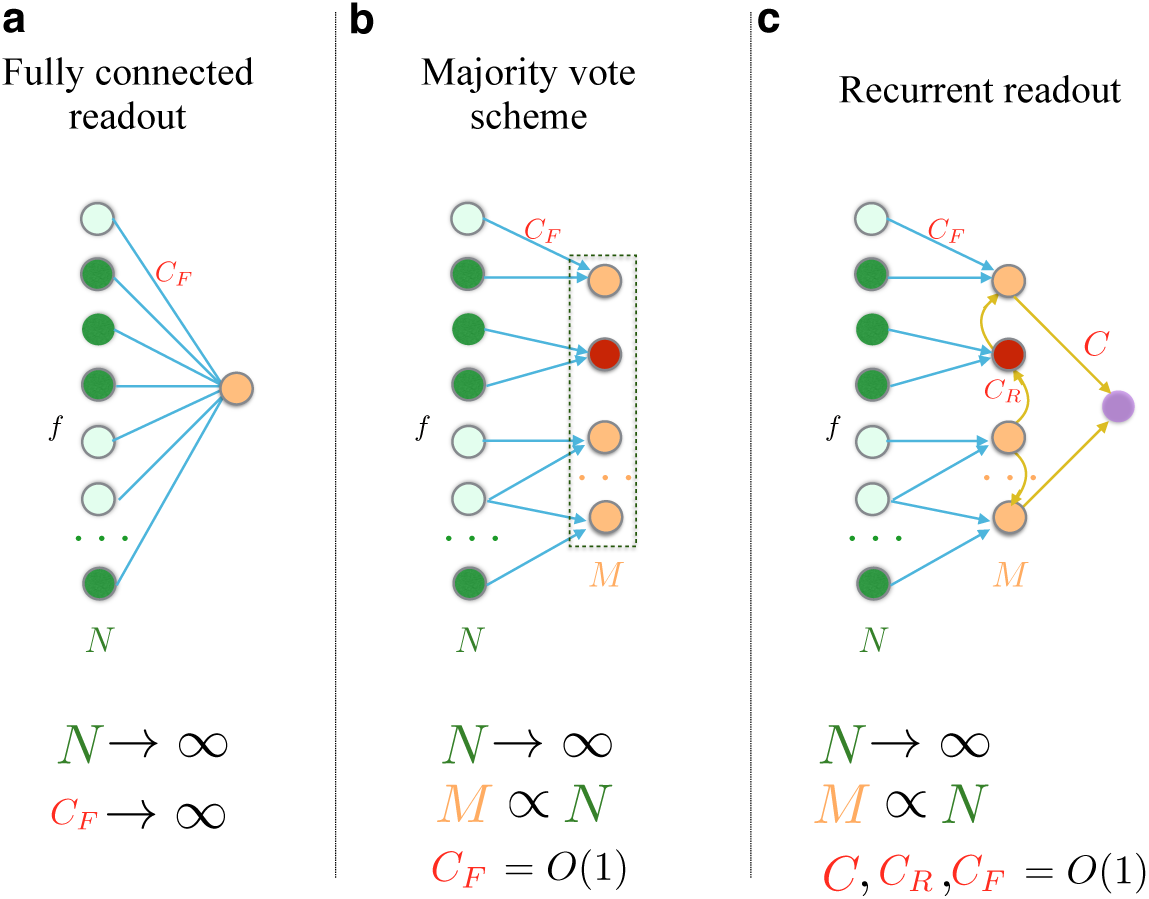
Architectures of the three network classifiers considered in the study, and their scaling properties. **a**. Fully connected readout, considered in section 3.1. The capacity of this classifier grows linearly with the number of input units *N*, however the number of afferent connections *C_F_* grows as quickly as *N*. **b**. Committee machine of partially connected perceptrons (section 3.2). The collective decision is made using a majority vote. Even though the number of connections per perceptron can be kept constant as the number of input neurons *N* increases, the number of readouts *M* has to grow with *N* in order to match the performance scaling of (a). The majority vote strategy requires another downstream readout, whose connectivity grows with *M* and hence with *N*. **c.** The recurrent readout that we propose in section 3.3. As *N* → ∞, the number of feedforward connections per perceptron *C_F_*, the number of recurrent connections per perceptron *C_R_*, as well as the number of connections of the downstream readout stay constant when *N* increases.

#### 3.1.1 Input statistics

We assume that pairs (*ξ^μ^*, *η^μ^*) of a pattern *ξ^μ^* and a label *η^μ^* are drawn from a random ensemble of *P* pairs (pattern, label). The pattern components 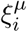 on all *N* input units and labels *η^μ^* are random independent variables. We assume that each component 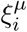 (*i* = 1… *N* is the unit index and *μ* = 1… *P* is the pattern index) is activated to 1 with probability *f* called *coding level* and otherwise is 0, and that label *η^μ^* takes one of the two values: *η^μ^* = +1 with probability *y*, called the output sparseness, and *η^μ^* = −1 otherwise:

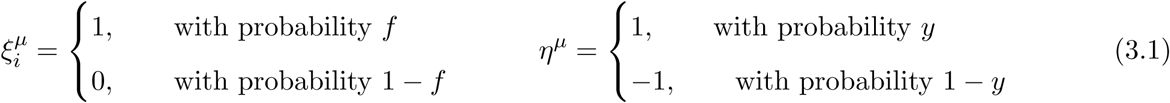

#### 3.1.2 Learning rule and the synaptic current

The linear threshold readout, or perceptron, classifies its inputs based on the sign of the weighted sum of the input components. This sum is sometimes called *synaptic current*, as it is viewed as modeling the synaptic current into a biological neuron

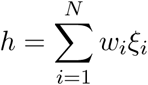

We say that the network has learned the association between *P* input patterns 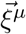 and *P* labels *η^μ^* if for any pattern *μ*

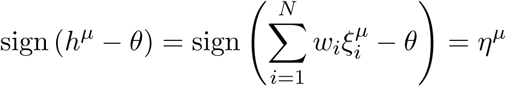

Where *θ* is the threshold, that we further assume to be equal to zero.

Training the network means finding the set of weights w_i_ that satisfies the above expression for all *P* patterns.

The Hebb-like learning rule, which we use to train the weights {*w_i_*} of the classifier is:

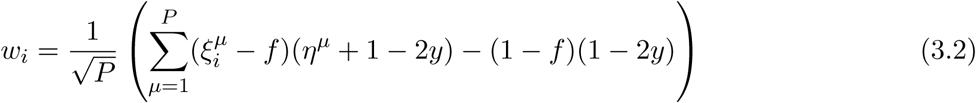

In the case when patterns are equally likely to belong to either class 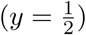, the learning rule simplifies to:

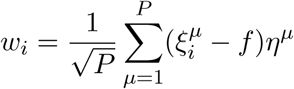

Here and in all that follows we set the threshold *θ* to zero.

After training, the synaptic current in response to a test pattern 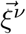 is

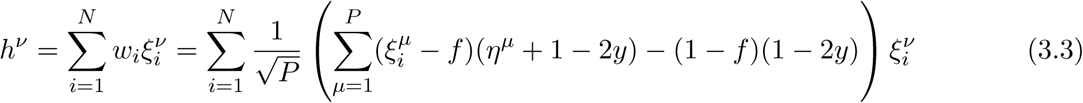

If 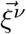 together with its label *η^v^* was part of the training set, we can split the sum over patterns into the contribution from the presented pattern 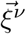 and the contribution from other learned patterns as follows

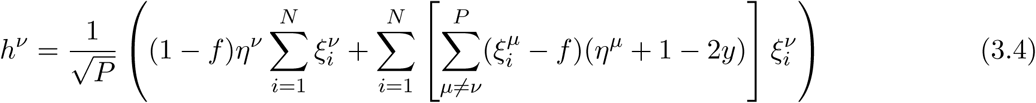

Here we used 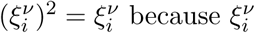 takes value 0 or 1.

We denote the number of active input units for the pattern *ν* by *n^ν^*

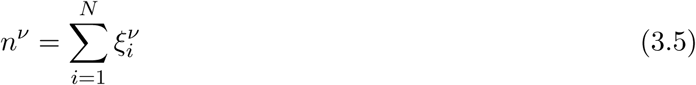

The value of *n^ν^* is in *binomial distribution* of *N* trials with probability *f*, **B**(*N*, *f*). Its expected value is determined by the number of input units *N* and the coding level *f*

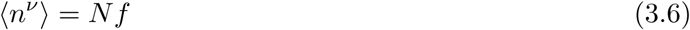

(here and throughout this text the angular brackets denote the mean over the realizations of the input patterns).

We replace the sum in the square brackets of (3.4) by 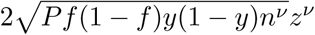 where we have introduced a *noise random variable z^ν^* with zero mean and unit variance. The coefficient is concluded from the fact that each individual term 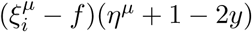 has variance

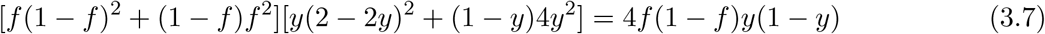

and the fact that the 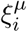 variables are mutually independent. By the central limit theorem the noise variable *z^ν^* can be approximated as Gaussian in the limit *P* → ∞ with finite *f* and *n^ν^*.

In terms of *z^ν^* and *n^ν^* the synaptic current is written as

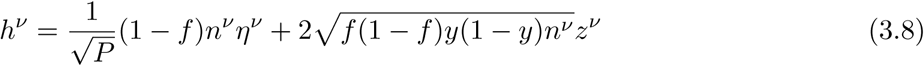

If a pattern belongs to either class with equal probability 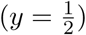, this expression simplifies to

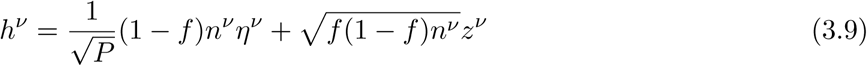

Note that the first term is the one that reflects the correct classification of the input pattern, and the second one represents the noise caused by the interference from other patterns that were learned by the perceptron. The important parameter is the ratio of the two, which is proportional to 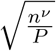

### 3.2 Committee machine

We now turn to deriving the classification capacity of a committee machine, the network shown on the Figure 1b, where each out of *M* perceptrons receives feedforward connections from *C_F_* input units. The connectivity *C_F_* does not scale when the number of input units *N* increases.

The final decision is the majority vote of the classifiers. In other words, if classification is accurate

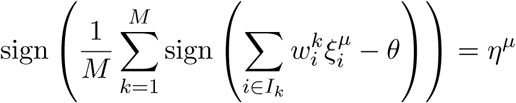

Here *i* ∊ *I_k_* stands for all the input units (there are *C_F_* of them) that are connected to the readout *k*, and 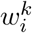 is the strength of the connection from the input unit *i* to the readout *k* (for the learning rule we consider 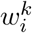 does not depend on *k*).
The synaptic current into the readout unit *k* when pattern 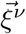 is presented is determined by

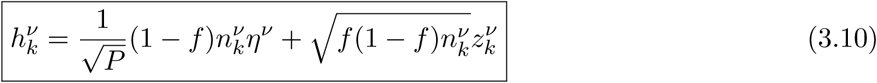

The number of active inputs connected to the perceptron *k*, 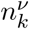 is drawn from the binomial distribution **B**(*C_F_*, *f*) of now *C_F_* trials with the success rate *f*: and its expectation value is

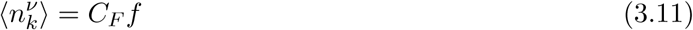

Since the number of connections per readout *C_F_* stays constant as the number of patterns *P* and the size of the network (*N* and *M*) grow, the probability of a single perceptron to classify a pattern correctly approaches the chance level. Indeed, in contrast to the fully connected perceptron, the number of active inputs 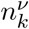 per readout neuron does not change with the size of the network (see (3.11)). Hence, the first term of the expression (3.10) decreases in the absolute value as the number of patterns *P* grows, while the typical value of the second term stays the same. However, there is always a slight tendency towards the correct answer 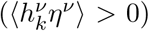, that can be utilized by having a growing number of sparsely connected classifiers that take a collective decision by majority vote. This scheme is known by the name of committee machine and has been shown to largely exceed the performance of a single classifier.

It is important to note that in order for the capacity of a committee machine to keep increasing as new classifiers (committee members) are added, the responses of different classifiers should stay sufficiently independent from each other. In the case of limited connectivity, which we consider here, the correlations automatically become smaller and smaller as we increase the number of input units. This happens because the probability of a typical pair of readout neurons to have a common input unit, and thus correlated responses, decreases. In order for the correlations not to be a limiting factor of the classification capacity, we need to increase the number of input units linearly with the number of perceptrons. If one introduces some other mechanism of reducing the correlations between the responses of the classifiers with common input units (like making different perceptrons learn different sets of patterns), a sublinear scaling of the number of input units *N* with the number of perceptrons *M* might be sufficient.

#### 3.2.1 Non-overlapping case

The majority vote of *M* linear threshold classifiers is given by the *average vote*

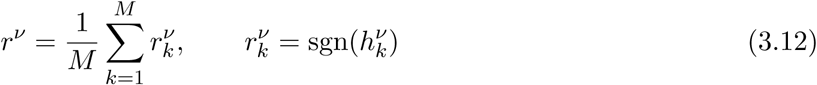

where 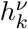 is given in (3.10). Positive *r^ν^η^ν^* means that the pattern ν is classified correctly.

The expectation value of *r^ν^* follows from (3.10) after integrating over the noise variable 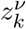, which is approximated to be normally distributed. We make an assumption 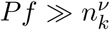, which is justified for a large number of patterns, and that allows us to use the approximation of the error function for small arguments to get

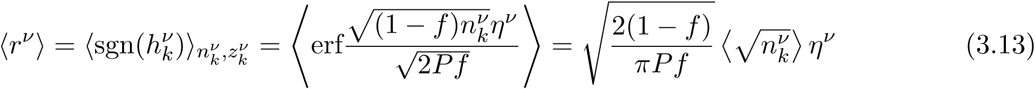

The expectation value 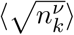 is computed over the binomial distribution **B**(*C_F_*, *f*)

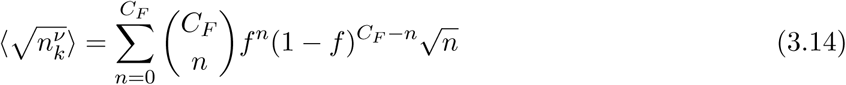

In the dense regime, *C_F_f* ≫ 1, it can be approximated by

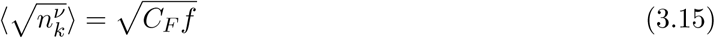

and in the extremely sparse case, when *C_F_f* ≪ 1 and only 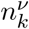 = {0, 1} are encountered substantially often, by

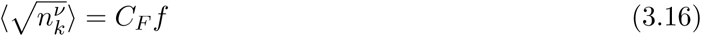

To proceed with deriving the classification capacity, let us start with independent classifiers first. The independence of the responses can be achieved either by forcing the connections to be non-overlapping, or by assuming an additional mechanism that, for example, causes different classifiers to update their incoming connections in response to different subsets of the input patterns.

In this case *r^ν^* can be though of as drawn from a Gaussian distribution with the mean given by (3.13) and the variance

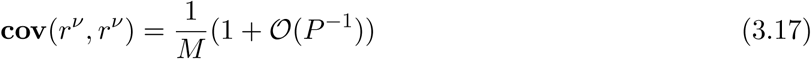

The gaussian assumption is justified by the law of large numbers.

Here and from now on we ignore the contributions of the subleading order, 𝓞(*P*^−1^) in this case. The probability *p*_Correct_ to classify a pattern correctly (*r^ν^η^ν^* > 0) can then be easily computed. Fixing *tolerated error rate* ε and requiring *p*_correct_ > 1 – ε leads to the expression for the maximal number of input patterns that can be classified with the accuracy 1 – ε.

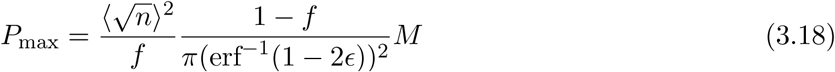

Here 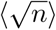 denotes the average over binomial distribution, *n* ∼ **B**(*C_F_, f*).

This result only holds for the case of non-overlapping connections or in the presence of a decorrelation mechanism. In the following section we generalize it to random connectivity.

#### 3.2.2 Correction to classification capacity due to overlap in the connections

To derive an analogous expression for the overlapping case without a decorrelation mechanism we need to compute the variance

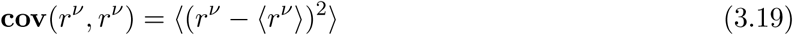

of the average vote *r^ν^*, defined by (3.12), taking into account the correlations of individual votes 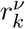.

We start by splitting the covariance into diagonal and non-diagonal contributions:

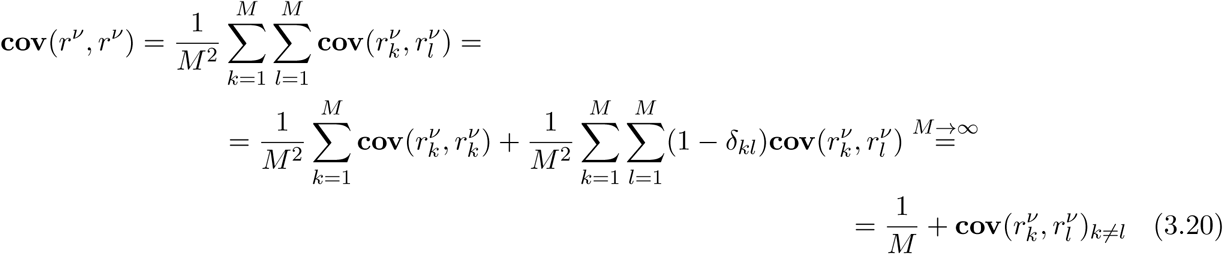

We assume that *M* and *N* scale linearly with *P* and *M*, *N*, *P* → ∞. The leading terms are thus of the order 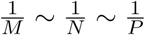 and we ignore all the subleading contributions.

When the classifiers *k* and *l* share input units, the correlation between their responses is positive and is closely related to the correlation of the input currents 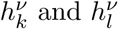 (see (3.10)).

Let 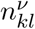 be the number of input units that are connected to both the classifier *k* and the classifier *l* and are active in the pattern 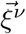. For a large number of input units *N* and finite connectivity Cf we can assume that 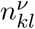 can be either 0 or 1, but not more. The probability of 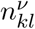 being 1 is given by

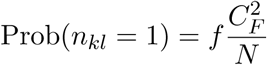

The number of active units that are connected to only one of the two classifiers are denoted by 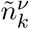 and 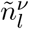 respectively. In the current approximation both of them can be assumed to be distributed according to a binomial distribution **B**(*C_F_*, *f*).

Then, the currents can be written as (see (3.10)):

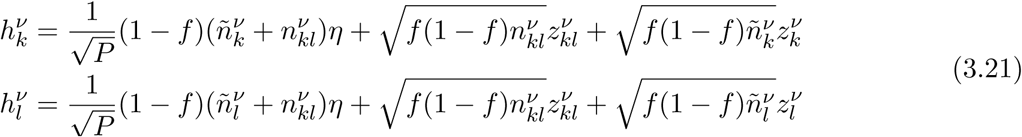

Where 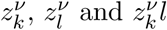 are all independent gaussian variables with zero mean and unit variance.

To compute the covariance

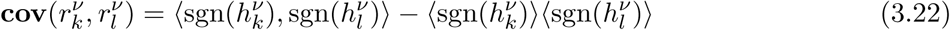

we start by integrating over the variables 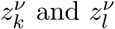 to get

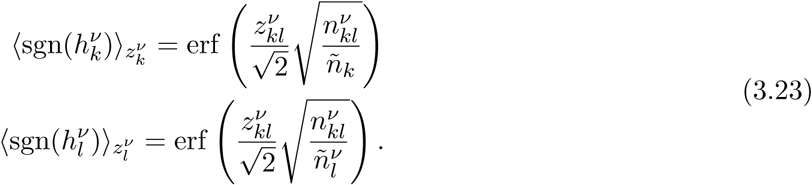

Then, (3.22) can be evaluated using the table integral ^1^

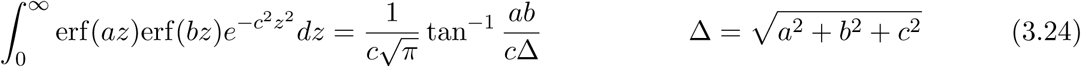

In the leading order we get:

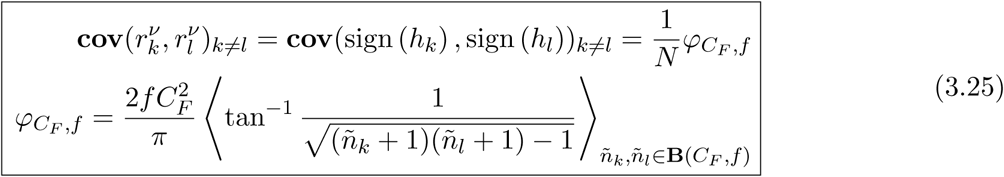

In the dense regime (*C_F_f* ≫ 1) the expression for *φC_F,f_* in (3.25) can be approximated as

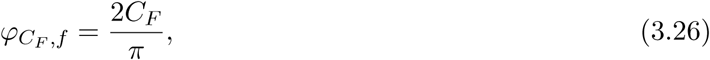

which leads

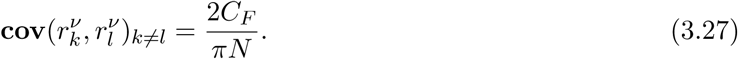

While in the sparse approximation (*C_F_f* ≪ 1),

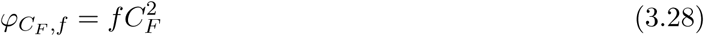

and

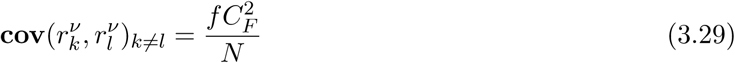

Plugging this result into (3.20), we get for the variance of the majority vote *r^ν^* in the overlapping case

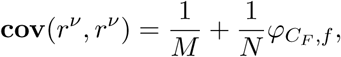

which together with (3.13) leads for the maximal number of input patterns that the committee machine can learn to classify with the accuracy 1 – ε

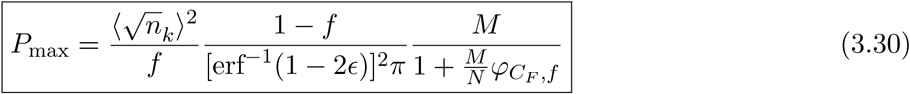

Here *φC_F,f_* is given in (3.25) and is approximated by (3.26) or (3.28).

If both the number of input units *N* and the number of classifiers *M* increase in proportion to each other, the capacity *P* increases linearly with *N* (or *M*).

In the case of dense representations, *C_F_f* ≫ 1 the last expression simplifies to

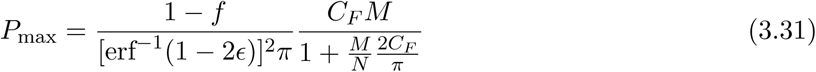

and in the ultra-sparse limit, *C_F_ f* ≪ 1

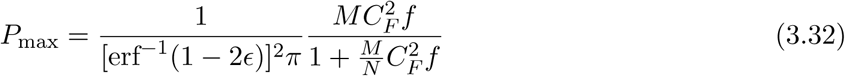

### 3.3 Committee machine with recurrent connections

The majority rule scenario already overcomes the limitations of the connectivity of a single perceptron, but this is not the final answer to constructing a classifier with limited connectivity. The reason is that we still need to implement the majority rule and bring the classification signal to the level of a single unit. The naive way to do it would require another final readout that would have to sample the entire population of *M* intermediate layer perceptrons. Since *M* has to scale linearly with the number of learned patterns *P*, the connectivity of the final readout would also have to scale linearly with *P* (see (3.30)) and would exceed any predetermined limit for sufficiently large number of learned patterns.

To implement the majority vote of the intermediate perceptrons while keeping the connectivity of any unit in the network limited, we introduce the recurrent connectivity in the layer of perceptrons. Our goal is to have two attractor states of the intermediate layer dynamics, that correspond to the two classes. The feedforward input through the connections {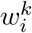} trained in the same way as before, will be slightly biased in the positive direction for one class of the input patterns and in the negative for the other. This slight bias determines which attractor state the network will choose. It is essential that the attractors are far away and do not become closer when the number of learned patterns *P* increases implies that the final readout will be able to discriminate between these states, and thus indicate the class of the presented pattern, even if its connectivity does not scale with *P*. It turns out that for binary classification it is enough to have random recurrent connectivity with sufficiently large but not increasing with *P* number of connections per unit. The weights of these recurrent connection do not have to be tuned (no learning required for recurrent connections).

We compute the probability of the network of recurrently connected readouts to go to the correct attractor (the one assigned to the class of the input pattern presented) as a function of the number of input units *N*, number of perceptrons *M* and various parameters of the network.

#### 3.3.1 Network topology

The recurrent readout network shown on the right of Figure 1c consists of the input layer (green), the intermediate layer of perceptrons (orange) and the final readout unit (purple).

As before, the *input layer* of *N* neurons is presented with a random and uncorrelated pattern 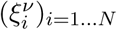 from a set of *P* patterns 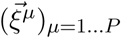 that the network has learned to classify.

The layer of perceptrons we now call *intermediate layer.* It consists of *M* linear threshold readouts, each of which is connected to a randomly chosen *C_F_* out of *N* input units. Hence, the *feedforward connectivity C_F_* is the number of feedforward inputs that each perceptron receives. The *C_F_* is an important parameter in the problem as it determines the classification capacity of a perceptron considered in isolation. The intermediate layer is recurrently connected. For the case of binary classification, the probability that two units are connected is the same for each pair. The recurrent connections are not plastic and can be chosen to be all of equal strength *α*.

The *final layer* consists of a single readout unit that is connected to a randomly chosen subset of *C* perceptrons in the intermediate layer, with the strength of all connections taken equal.

The recurrent connectivity matrix *J_kl_*, *k, l* ∊ [1… *M*] is assumed to be symmetric

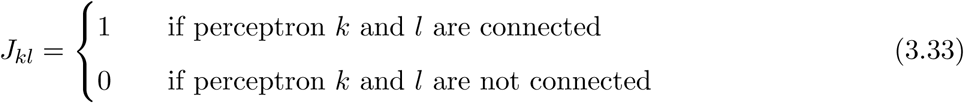

Let *C_R_* be the number of recurrent connections per unit

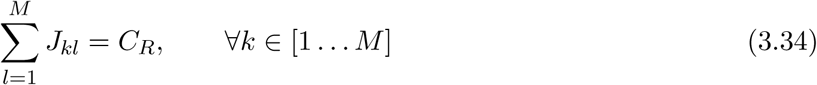

We will keep the connectivity parameters *C_F_*, *C_R_* and *C* and coding level *f* at fixed constant values, while sending the number of input units *N*, the number of intermediate perceptrons *M* and the number of patterns *P* to infinity

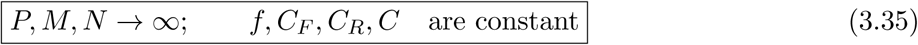

We want to recover the linear scaling of the maximal number of patterns *P*_max_ that the network can learn to classify with the number of input units *N*, which is known to hold for the fully connected perceptron [28].

#### 3.3.2 Discrete time dynamical model

We model the recurrent dynamics as a probabilistic dynamical process in discrete time *t* with the probabilistic transition rule from a network state at time *t* to a network state at time *t* + 1. Let *s_k_*(*t*) ∊ [−1, 1] for *k* ∊ [1… *M*] be the dynamical variable describing the state of unit *k* at time *t* in recurrent network.

Let 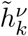 be the total current into the readout unit *k*

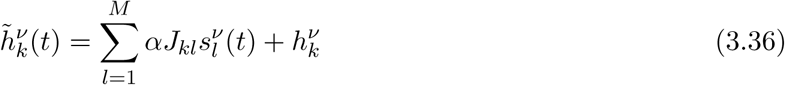

where the first term corresponds to the recurrent contribution and the second term represents the feedforward current from the input layer (3.10) that is constant in time.

The probabilistic transition rule from the state at time *t* to the state at time *t* + 1 is

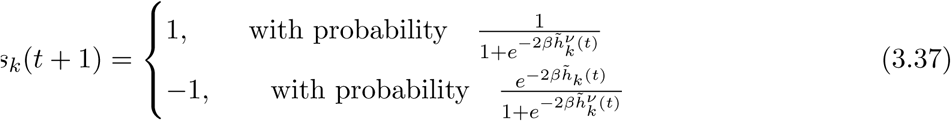

Here *β* is the *inverse temperature parameter* for the statistical model of the recurrent dynamics. We approximate this probabilistic recurrent dynamics with the *mean field* method.

#### 3.3.3 Mean field analysis of the recurrent dynamics

To compute the capacity of such a recurrent classifier, we analyze the recurrent dynamics in the mean field approximation. The activities of the recurrently connected units are represented by the variables *s_k_* = {+1, −1} with *k* = 1… *M*. The mean field equation for the *average activation* of the recurrently connected intermediate layer in response to the pattern *ν*,

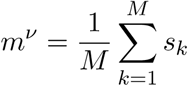

reads

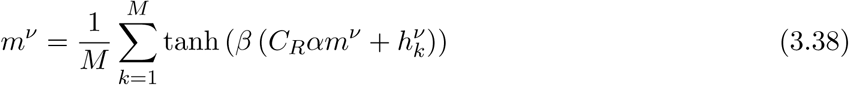

The average activation *m^ν^* is close to zero if the amount of active and inactive units is approximately the same. If the majority of the units is in the active state, *m^ν^* will be close to 1, and if the majority is inactive, *m^ν^* will be close to -1.

Here *C_R_* is the average number of connections per unit, *α* is the strength of recurrent synapses (we assume they are all excitatory and of equal strength), *β* is the inverse temperature parameter and 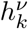 is the feedforward input current given by (3.10).

We proceed by analyzing the above equation graphically. The plot of the right-hand side is a sigmoid curve and the left-hand side is a line at 45 degrees. The intersections of these two lines determine the solutions to the equation. There are two possible situations that correspond to two different scenarios of the network dynamics.

The first scenario, shown on the Figure 2a and 2b, is characterized by having only one point of intersection of the line and the sigmoid. In this case there is only one solution to the mean field equation (3.38) and only one stable state of the recurrent network. The right hand side of the equation is almost but not quite an odd function of its argument *m^ν^*, so the sigmoidal curve representing it is slightly shifted to the left if 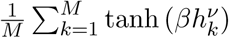 > 0 and to the right if 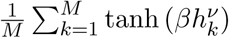 < 0. If the curve is shifted to the left, the single point of its intersection with the strait line passing thorough the origin will be in the right half-plane. So, for the positive input pattern (*η^ν^* = + 1 and 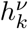 is more likely to be positive) the mean activity of the intermediate layer in the stable state *m^ν^* will usually be positive, while for the negative input patterns it will be negative. Even though there is a relation between the sign of the mean activity of the intermediate layer in the stable state and the class of the input pattern, this is not helpful for our purposes. The reason is that we encounter exactly the same problem as for the case of no recurrent connections: the absolute value of the average activity *m^ν^* will decrease with the number of learned patterns *P*, which means that the number of active and inactive units in the intermediate layer will become more and more similar. Consequently, to sample this small imbalance we would require larger and larger connectivity of the final readout. In short, the regime with one stable solution (Figure 2a and 2b) is not much different from the case of no recurrent connections. Not surprisingly, this regime corresponds to relatively weak recurrent connections.

**Figure 2.**
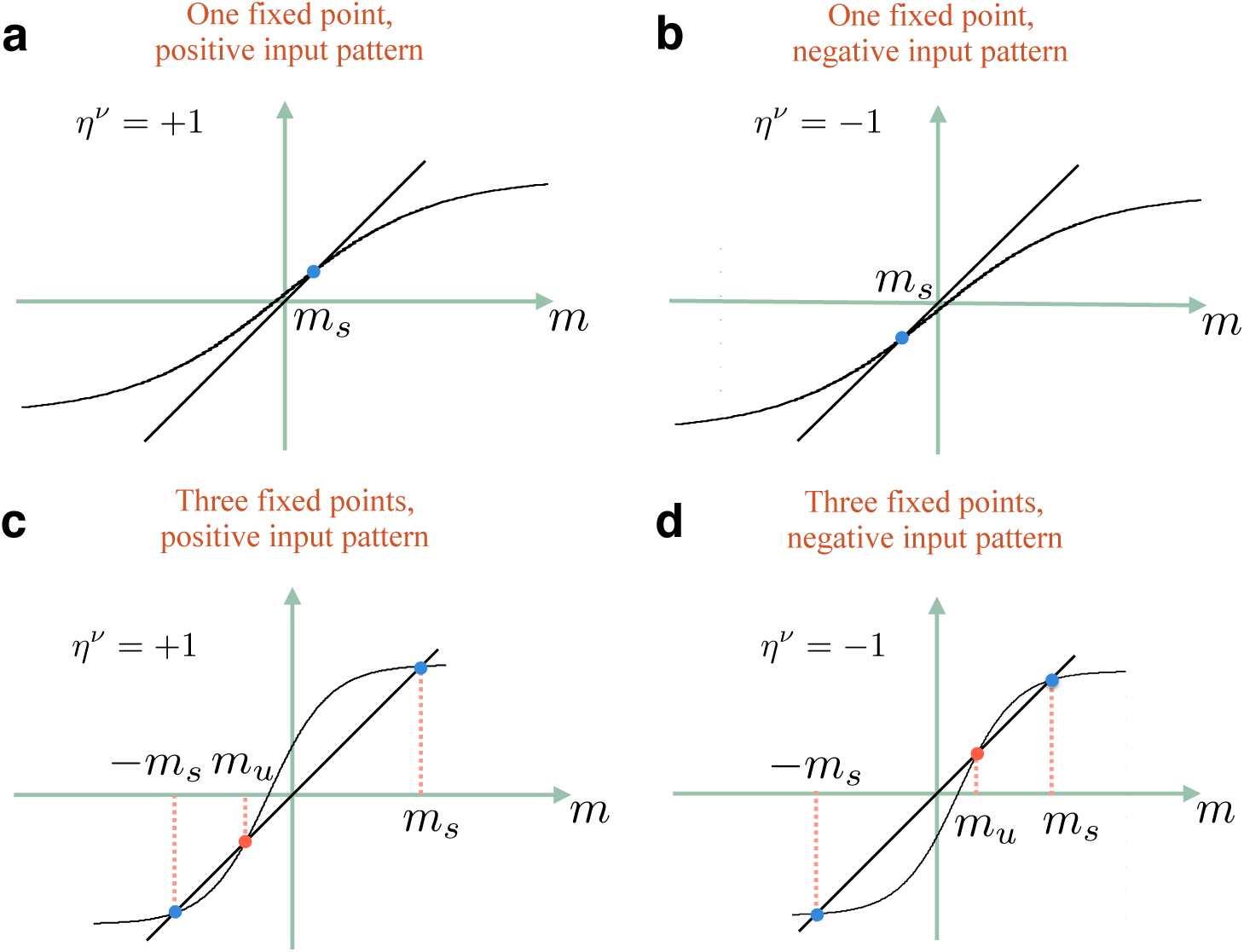
Graphical representation of the mean field equation (3.38). The left-hand side of the equation is represented by the line, and the right-hand side - by the sigmoidal curve. The slope of the sigmoidal curve is determined by the amount of dynamical noise in the update rule relative to the strength of the feedforward connections (*βC_R_α*), and the shift relative to *m* = 0 is based on the expected value of the feedforward input 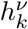 **a**. When the input pattern belongs to the “positive” class and the noise is high, there is only one solutions to the equation, which corresponds to small but positive value *m* = *m_s_*. This solutions is stable. **b**. For “negative” input pattern, the solution is negative, *m_s_* < 0. **c,d**. In the case of low noise, there are three solutions to the mean field equation, with tow extreme solutions *m_s_* and −*m_S_* being stable, and the middle one *m_u_*, which is close to zero, being unstable. For the case of positive input pattern *m_u_* < 0, and for the case of negative pattern *m_u_* > 0.

It is the other situation, shown on the Figure 2c and 2d, that is actually of interest. There are three points of intersection of the sigmoid curve of the right hand side of the equation (3.38) and the straight line of the left hand side. The stable states of the network correspond to the rightmost and the leftmost solutions, that are both characterized by a large imbalance between active and inactive units (|*m^ν^*| ∼ 1). Most importantly, these solutions are virtually insensitive to the distribution of 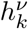, and hence to the number of learned patterns *P*. So, if we postulate that the right solution corresponds to the positive input patterns and the left solution to the negative ones, it will be easy for a downstream readout with connectivity that does not increase with *P* to distinguish between them.

The middle intersection point *m_u_* corresponds to the unstable solution. When the network is initialized at the state 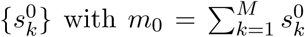 on the left of the unstable solution *m*_0_ < *m_u_*, the recurrent dynamics will most likely evolve to the left stable state, and if initialized at *m*_0_ > *m_u_* it will evolve to the right stable state. As shown on the Figure 2c,d the point of unstable equilibrium will be to the left of the origin for a positive input pattern and to the right of the origin otherwise (due to the difference in the mean of the distributions of 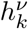). Hence, initiating the network at *m*_0_ = 0 will serve the purpose of biasing the evolution of the network towards the stable state that corresponds to the class of the input pattern. If the number of learned patterns *P* is large, the point of unstable equilibrium is very close to zero 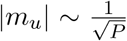, this is the manifestation of the same problem as before, namely the decrease of the signal to noise ratio with the increasing number of learned patterns. Thus, the noise in the initial state of the network *m*_0_ should also decrease as 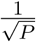. This is achieved if all the units in the intermediate layer are initialized at 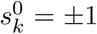 with equal probabilities independently from each other, and the number of units *M* is linear in *P* (the same scaling as for the committee machine discussed earlier). We use this initialization process to derive the classification capacity and to run the simulations. In the section 4.6 we suggest a biologically plausible way to initialize the network at the desired point.

To summarize, the information about the class of the input pattern is contained in the feedforward input to the intermediate recurrently connected layer. In the case of a single stable state (Figure 2a and 2b), although average activity of the network reflects this information, the signal is very small and a fully connected downstream readout is required. In the case of two stable states (Figure 2c and 2d), this small signal biases the network to choose the one corresponding to the class of the input pattern, and by doing so, the network amplifies the feedforward signal making it easy to read out by a sparsely connected downstream unit.

#### 3.3.4 Number of classifiable inputs

As discussed in the previous section, the requirement for the correct classification of an input pattern by means of recurrently connected committee machine is that the average activity of the network at the initial moment 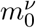 is on the correct side of the point of unstable equilibrium 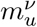, namely

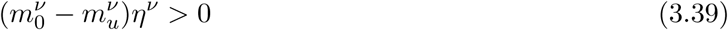

where *η^ν^* is the desired output (*η^ν^* = {±1}).

In what follows we drop the pattern index *ν*.

The statistics of *m*_0_ over random initializations of the network follows from its definition

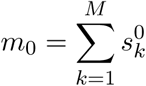

where each unit is initialized at *s_k_* = +1 or *s_k_* = −1 with equal probability:

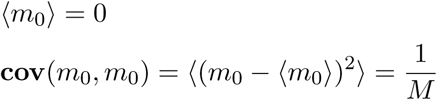

Since *M* is a large number, we approximate the distribution of *m*_0_ by a Gaussian distribution with these mean and variance.

The position of the unstable equilibrium point *m_u_*, corresponding to one of the three solutions (the one that is close to zero) of the mean field equation (3.38), can not be computed analytically in the general case. However, there are parameter regimes in which we can compute the approximate first and second order statistics of *m_u_* over random realizations of the input patterns. These parameter regimes and corresponding approximations are discussed in the following section. Once the mean *μ_u_* which depends on the number of learned patterns *P*, and the variance 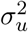 of *m_u_* are known, the requirement to classify *P* input patterns with accuracy 1 ‒ ε can be written as (assuming the distribution of *m_u_* to be also Gaussian)

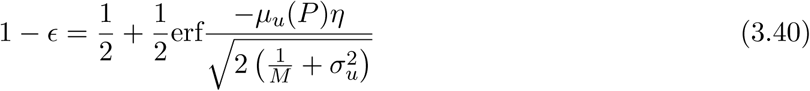

The expected number *P* of correctly classified patterns can be found by inverting the above equation.
In the following sections we consider different parameter regimes that lead to different approximations for *μ_u_* and *σ_u_*.

#### 3.3.5 The uniform regime

In the current study, among other issues we are interested in the consequences of the sparsity of input representations. Since we consider the feedforward connectivity *C_F_* to be a constant number and not to scale with the size of the network, for sparse representations there will be a substantial number of perceptrons that receive zero feedforward input. Unless the dynamical noise is very high, these units should be considered separately, and in the mean field approximation an additional order parameter should be introduced to describe their average activity. We call these units *free units*.

Uniform regime is the parameter regime under which it is not necessary to analyze the free units separately, and the equation (3.38) is valid without modifications. Obviously, when the input representations are dense, *C_F_ f* ≫ 1, the network of the intermediate layer is in the uniform regime, since there are not enough of free units to make a difference. However, assuming the uniform regime is also valid independently of the number of free units, when the dynamical noise is very large in comparison with the feedforward input (see the next section).

The conditions defining the uniform regime are:

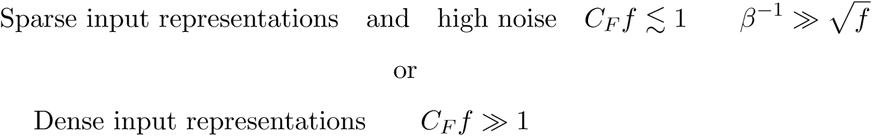

#### 3.3.6 Uniform regime, high noise

One approximation we can make to find the unstable solution *m_u_* of the mean field equation (3.38) is the high noise approximation, which is defined by the requirement

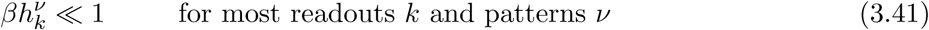

It follows from the expression (3.10) for the feedforward current that this requirement is met if

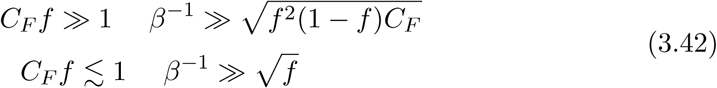

The condition for having three solutions of equation (3.38) rather than one (see Figure 2) is

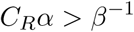

Since we are looking for the solution, which is close to zero and (3.41) is satisfied for most of the terms, the equation (3.38) can be approximated by replacing the hyperbolic tangent by its argument:

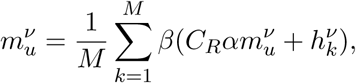

(note that this approximation is also valid for the terms with 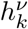 = 0).

Solving this equation leads for the mean *μ_ν_* and the standard deviation *σ_u_* of *m_u_*:

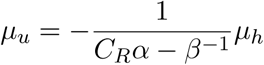

and

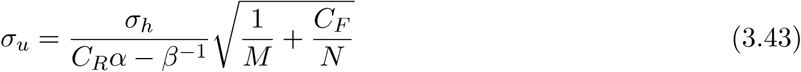

The mean *μ_h_* and the standard deviation *σ_h_* of the feedforward current 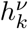 are computed from (3.10):

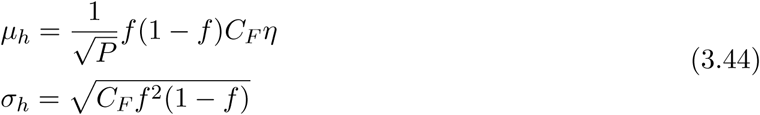

The *C_F_/N* term in (3.43) comes from the correlations between the feedforward currents 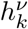 into different readouts *k* due to overlapping connections (see Appendix 6.1).

Now the maximum number of learned patterns for the classifier in the uniform regime for high noise approximation can be computed from (3.40) and is given by

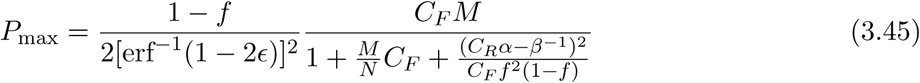

We note, that because of the applicability conditions (3.42) making the last term in the denominator small requires fine tuning of the parameter *β*.

#### 3.3.7 Uniform regime, low noise

The other approximation in which the equation (3.38) can be solved is

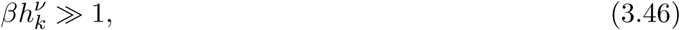

which is true if

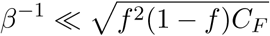

Under this condition, assuming the uniform regime is only valid if the input representations are dense

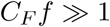

The condition for having three solutions to the mean field equation in the low noise approximation becomes (see (3.50))

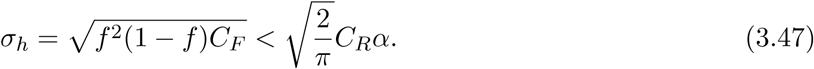

In this case the hyperbolic tangent in the equation (3.38) can be approximated by the sign function

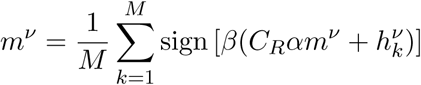

Let us denote the right side of this equation by g(*m_u_*), where

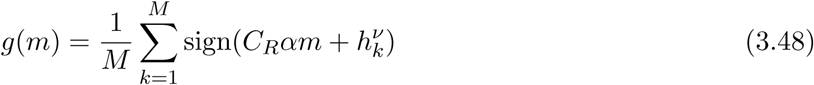

is a stochastic function over different realizations of {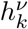}.

Note that in this case, having a substantial fraction of terms with 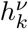 = 0 would lead to a discontinuity of the right hand side at 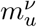 = 0.

The mean 〈*g*(*m*)〉 can be found by integrating over the distribution of 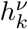 (see (3.10))

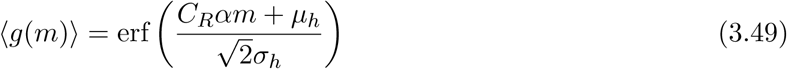

Where *μ_h_* and *σ_h_* are the mean and standard deviation of 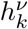 respectively, which are given by (3.44).

Thus, when averaged over training patterns, the mean field equation becomes

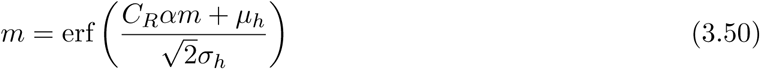

and it has three solutions when the derivative of the right-hand side with respect to *m* at *m* = 0 is larger than 1, which for *μ*_*h*_ ⋘ *σ_h_* immediately leads (3.47).

We now return to estimating the mean and the standard deviation of *m_u_*, which is the unstable solution to the approximated mean field equation

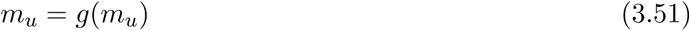

where *g*(*m*) is defined by (3.48).

For *μ_h_* ⋘ *σ_h_*, which is always the case if the number of stored patterns *P* is large enough, we assume that *C_R_αm_u_* is also small compared to *σ_h_* and check the self-consistency later. Then, we can use the approximation for the error function at small arguments to get

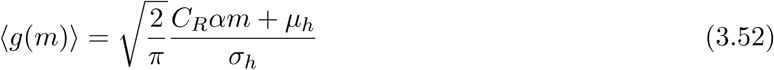

the variance of *g*(*m*) can be written as as sum of the diagonal and the non-diagonal terms

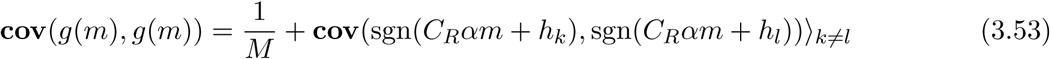

which is similar to the expression (3.20) for the variance of 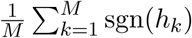 computed previously in (3.25). The only difference is that here the distribution of *h_k_* is shifted by *C_R_αm*. However, because the mean 〈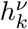〉 did not affect the result (3.25) and *C_R_αm_u_* + *μ_h_* is still negligible compared to *σ*_h_, we can write

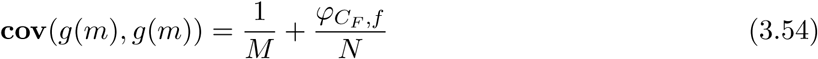

where *φC_F_,f* is given in (3.25).

As a sum of large number *M* of weakly correlated terms, *g*(*m*) can be assumed to be normally distributed and can be written as

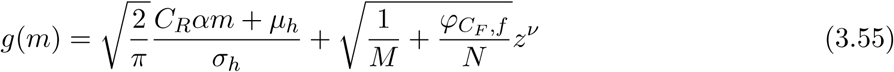

Where *z^ν^*is a Gaussian variable with zero mean and unit variance.

Plugging the expression for *g*(*m*) into (3.51), and solving for *m_u_* we get

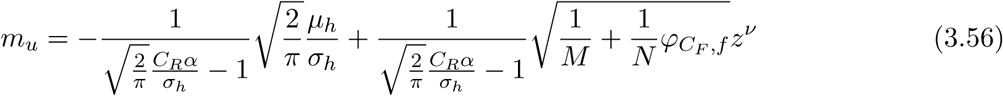

where *φC_F_,f* is given in (3.25).

So, the expectation value of *m_u_* is

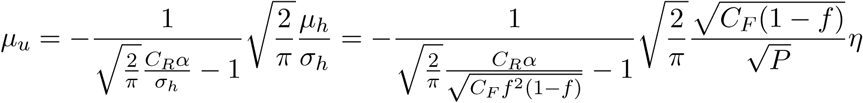

and the standard deviation is given by:

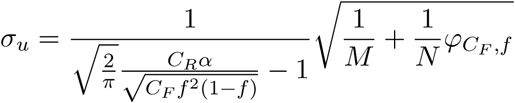

Because uniform regime and low noise implies dense input representation, we can use the dense approximation (3.26) for *φC_F_,f*. Plugging these results into (3.40) leads the capacity for the uniform regime, low noise

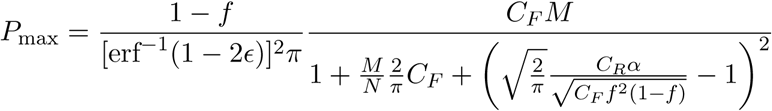

#### 3.3.8 Non-uniform regimes

When the input representation is sparse

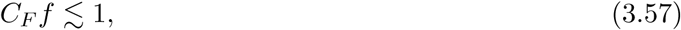

there is a substantial fraction of perceptrons for which all inputs are silent, we call them the free units. If the noise is not very high 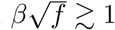 1, these units are statistically different from those that do receive a non-zero input. To analyze such a system in the mean-field approximation, two order parameters and two coupled mean-field equations should be introduced. To avoid this complication we consider a simpler case, to which we refer to as the *two-subnetworks* regime. This regime is characterized by the recurrent connections that are relatively weak when compared to the feedforward ones, so that the state of those units that do receive non-zero feedforward input is determined by this input only. Neither recurrent input nor noise can flip them. Only the free units participate in the recurrent dynamics and their mean activity in the final state reflects the class of the input pattern. Which of the two stable states the subnetwork of free units will go to is biased by the input from the input receiving units, which do have the information about the class of the input pattern from the feedforward input.

This approximation is valid if

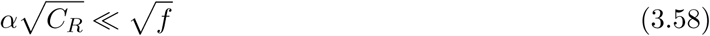

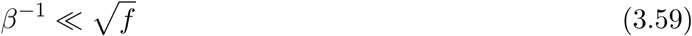

To be more precise, the former condition does not guarantee that the recurrent input will not be able to flip the input receiving units close to the final state, when most of the free units are synchronized. However, if this is the case, their activity already reflects the correct classification of the input pattern, and the input receiving units will flip in the right direction.

The mean field equation (3.38) should now be seen as describing the subnetwork of free units, and should be modified in several ways.

First, the number of units in the network is

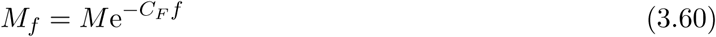

(since for small *f* the probability of all *C_F_* independent inputs to be silent is (1 – *f*)^*C_F_*^ ≈ e^−*C_F_ f*^). Second, only *C_R_*e^−*C_F_f*^ out of *C_R_* recurrent connections per unit come from other free units. Also, the external input to the network now comes from other (input receiving) units in the intermediate layer, rather than from the input layer.

The modified mean-field equation reads:

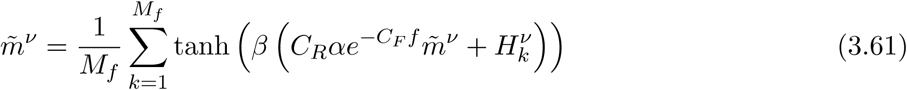

where 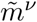 is the average activity of the subnetwork of free units and the index *k* runs over all the free units.

The external input

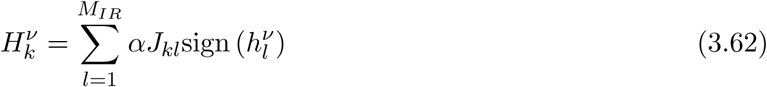

where the summation is over the input receiving units and 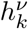 is the feedforward current of (3.10) with 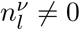. *M_IR_* is the number of input receiving units

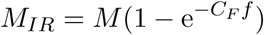

On average, the free unit *k* receives *C_R_* inputs, and (1 – *e^−C_F_f^*)*C_R_* of them come from input receivers. So (3.62) will have on average *C_R_*(1 – *e^−C_F_f^*) non-zero terms. Assuming that this is a large number, 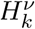 is a Gaussian variable with the mean given (in the leading order) by

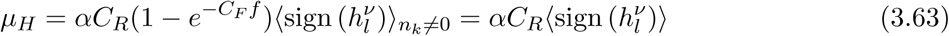

which using (3.13) becomes

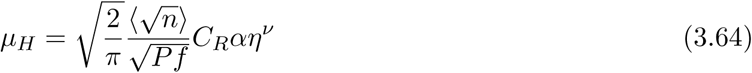

(assuming 1 – *f* ≈ 1). The number of active inputs *N* connected to the intermediate unit comes from the binomial distribution, *n* ∼ **B**(*N*, *f*).

The standard deviation of 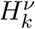 is

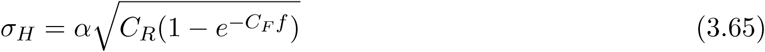

(the corrections due to correlations between different input receiving units are suppressed as 1/>*N* and will become negligible for large networks when *C_R_* does not scale with *N*).

To find the statistics of *m̃_u_*, the point of unstable equilibrium, we again consider high and low noise approximations, but now we should compare the inverse temperature parameter *β* to the standard deviation of 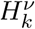.

What we further call *intermediate noise* is the noise which is small on the scale of the feedforward input (3.59) but large when compared to the typical values of 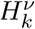.

#### 3.3.9 Two-subnetworks regime, intermediate noise

The following analysis is valid if in addition to the conditions (3.57), (3.58) and (3.59) the dynamical noise is high in comparison to the typical external input to the subnetwork of free units:

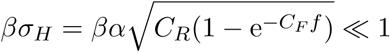

The condition for three solutions to the mean field equation (3.61) in this case is

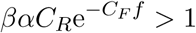

The former inequality allows us to approximate the hyperbolic tangent in (3.61) by its argument when looking for the unstable solution *m̃_u_*, which is close to zero:

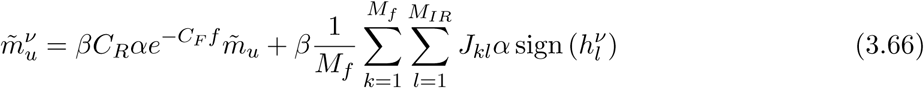

Each input receiving unit *l* has *C_R_* outgoing connections and approximately *e^-C_F_f^ C_R_* of them terminate on a free unit. Hence, the double sum can be rewritten as

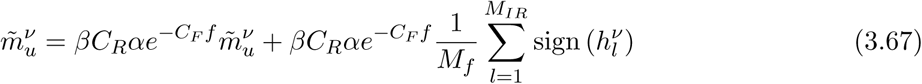

Solving this equation for *m̃_u_* leads (see (3.60))

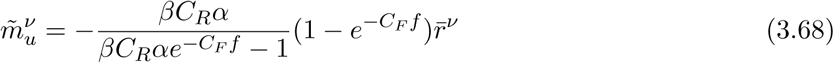

where we have introduced 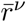: the sign of the feedforward current averaged over the units for which this current is non-zero

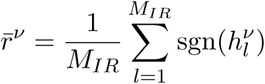

The statistics of 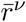 is closely related to previously computed statistics of *r^ν^* (see (3.12)), which is the sign of the feedforward current averaged over all the intermediate units. Namely,

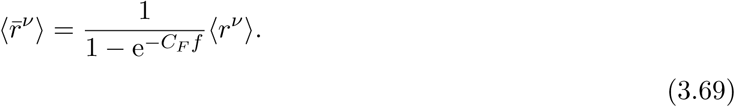

The expression for 〈*r^ν^*〉 is given in (3.13), which leads (we approximate 1 – *f* ≈ 1)

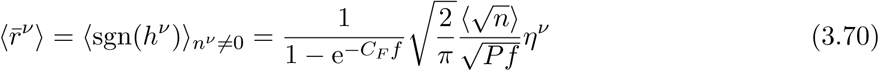

To compute the second order statistics of 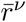, we use the relation

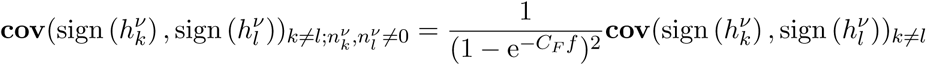

The covariance on the right-hand side was also computed in (3.25), which allows us to write

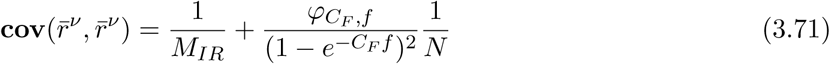

Plugging in (3.70) and (3.71) to (3.68) leads the expressions for the mean and the standard deviation of 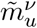:

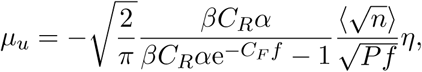

(the mean 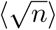 is computed assuming a binomial distribution for the number of active inputs *N* connected to a readout *n* ∼ **B**(*N*, *f*)), and

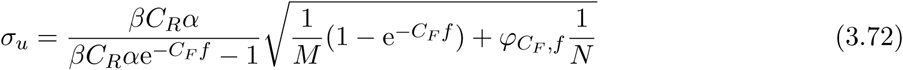

Now we can use (3.40) to compute the maximum number of learned patterns in the two-subnetworks regime under intermediate noise. The number of units in the network *M* in (3.40) should be replaced by the number of free units *M*e^*−C_F_f*^. The result is

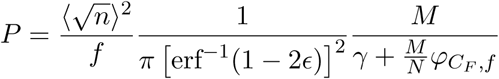

where

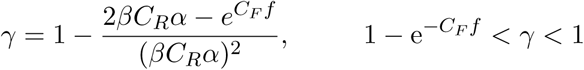

It is helpful for analyzing this result to rewrite the expression for *γ* in terms of Δ = e*^−C_F_f^ βC_R_α* – 1, which is the measure of how far the current parameters are from the transition to the one solution scenario (Figure 2a and 2b), at which the current framework breaks down.

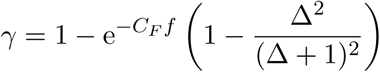

In the ultra-sparse approximation

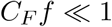

we can use (3.16) and (3.28) to get

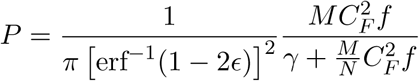

#### 3.3.10 Two-subnetworks regime, low noise

We now consider the low noise approximation to the mean field equation for the subnetwork of free units (3.61). This approximation is valid when in addition to (3.57), (3.58) and (3.59)

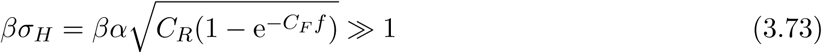

In this approximation, the mean field equation has three solutions if

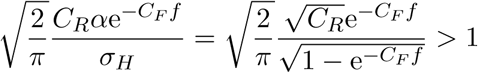

This condition is derived analogously to (3.46).

Under the assumption (3.73), the mean field equation (3.61) can then be approximated as

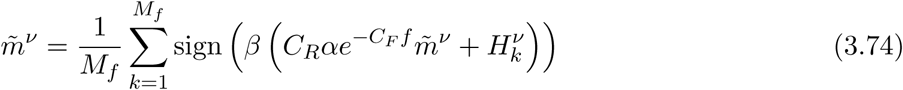

As in the section 3.3.7, let us introduce a stochastic function *g*(*m̃*)

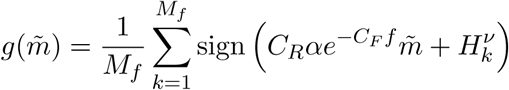

For small values of the argument *m̃*, the mean of *g*(*m̃*) over different realizations of 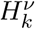 is approximated as

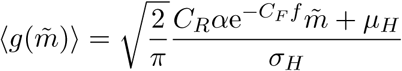

where *μ_H_* and *σ_H_* are given by (3.64) and (3.65).

To compute the variance of *g*(*m̃*) we need to know

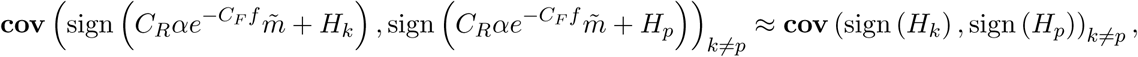

which is calculated in the Appendix 6.2, and for large absolute values of the recurrent connectivity (*C_R_e^−C_F_f^* ≫ 1) is approximated by

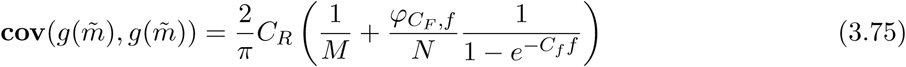

Assuming 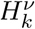 to be Gaussian, we can write

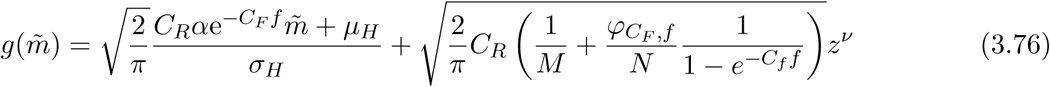

where *z^ν^* is a Gaussian variable with zero mean and unit variance.

The statistics of the unstable, close to zero, solution of (3.74) can now be found by plugging in (3.76) as the right hand side of (3.74), and solving for *m̃^ν^*.

After substituting (3.64) and (3.65) for *μ_H_* and *σ_H_*, we get for the mean and the variance of the unstable solution *m̃_u_* (assuming 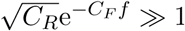):

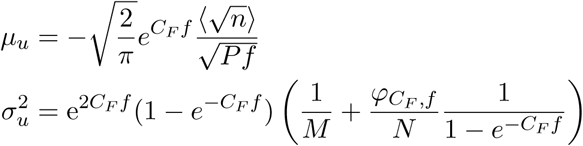

Using these expressions and (3.40) with *M* replaced by the number of free units *M_f_* = *Me^−C_F_f^*, we get for the maximal number of classifiable inputs in the low noise approximation of the two-subnetworks regime:

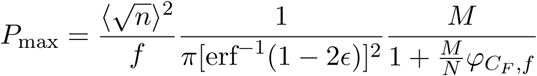

Note, that this is the same expression as (3.30) for the majority vote scenario (see Results section for an intuitive explanation).

For very sparse representations

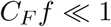

the expression simplifies to

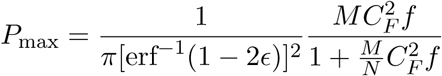

## 4 Results

### 4.1 The task and the network architecture

To evaluate the performance of different network architectures we consider a task in which the neural network is trained to associate a specific response to each input. The response is expressed by the activity of one output neuron, which could represent a decision, the expected value of an input stimulus or an action. Each input, for example a sensory stimulus, is a pattern of activity across *N* input neurons. Both, input and output neurons, are either active or inactive and hence the variables representing their activity are binary. Moreover, we assume that the inputs and the outputs are random and uncorrelated. Input neurons are active with probability *f*, whereas the output neuron is active on average for half of the inputs. Performing this task is equivalent to solving a binary classification problem in which each input is assigned to belong to one of two possible classes. As a measure of the performance of the network we introduce the classification capacity, the maximum number of input patterns that can be correctly classified, and determine how it scales with the total number of neurons in the network. We now consider architectures with increasing complexity and we eventually show that it is possible to design a network in which the number of classifiable inputs is large and it scales linearly with the number of neurons while each neuron has limited connectivity (i.e. the number of connections is fixed in the sense that it does not have to scale with the number of neurons).

#### 4.1.1 Single readout

The most basic network that we consider is the one in which the input neurons are directly connected to the output, which is basically the classical perceptron [1] (see Figure 1a). The network is trained by modifying the weights *w_i_*. that connect each input neuron *i* to the output. The output activity o^*μ*^ in response to stimulus *μ* is determined by thresholding the weighted sum of the inputs:

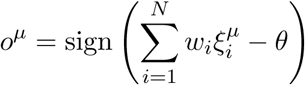

where *θ* is a threshold and 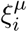 is the activity of neuron *i* when input pattern *μ* is selected. The weights *w*. and the threshold *θ* are learned to impose that *o^μ^* = *η^μ^*, where *η^μ^* is the desired output in response to stimulus *μ*. We know from many studies (see e.g. [28, 30]) that the maximum number of random inputs that can be correctly classified scales linearly with the number of input units when *f* = 1/2. This is a very favorable scaling, and actually the optimal one in the benchmark that we consider. Unfortunately, the number of connections of the output neuron is equal to the number of input neurons, and hence when the number of classifiable inputs grows, also the connectivity has to increase accordingly. This is true also in the case of sparse input representations. Indeed, for an arbitrary *f*, when we used a simple learning rule inspired by [31]

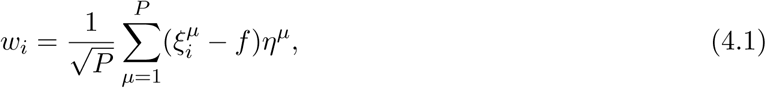

we obtained in the limit of large number of input neurons *N*, that the maximum number of input patterns that can be classified *P*_max_ is given by

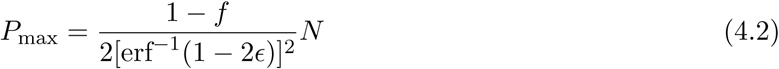

where ε is the maximum tolerated error.

Notice that the factor containing the coding level of the patterns *f* cannot change the scaling properties of *P*, even in the case in which the inputs become very sparse (i.e. when *f* → 0 as 1/*N*). This seems to be in contradiction with the results of [31, 32] in which *P* can scale as *N*^2^ when the inputs are sparse. However, it is important to remind that the *N*^2^ scaling can be achieved only when both the input and the output are sparse and in the cases that we analyzed here the output is dense (i.e. active in half of the cases).

We now consider a different architecture that partially overcome the limitation imposed by the limited connectivty assumption.

#### 4.1.2 Committee machines

Consider now the architecture of Figure 1b in which multiple perceptrons are combined together. We assume that each perceptron has limited connectivity, or more precisely, that when the number of input neurons becomes large (mathematically we consider the limit for *N* → ∞), the number of input connections per perceptron, *C_F_*, does not increase (i.e. remains finite when *N* → ∞). As a consequence, each perceptron will sample only a small fraction of the input neurons, and for this reason, it will misclassify most of the inputs when *P* becomes large (*P* → ∞). More quantitatively, the fraction of correctly classified inputs will be slightly above chance level (1/2), approximately 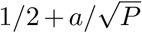 when *P* is large, *a* is a constant.

In this situation, each perceptron is said to be a weak classifier. However, if the responses of different perceptrons are sufficiently independent they can be combined together to perform significantly better than any individual perceptron. Multiple perceptrons combined together make what is called a committee machine. Typically the class of an input is decided by the committee using a majority vote rule: if the majority of perceptrons are active then the output neuron should also be active, otherwise it should be inactive. The majority rule can be easily implemented by summing with equal weights the outputs of all perceptrons.

As mentioned in section 2, adding new readouts without increasing the number of input units *N* can not increase the classification capacity indefinitely, unless an additional mechanism is introduced to decorrelate the responses of different readouts. Such mechanisms may very well exist in the real brain. For example, one could imagine some local changes of synaptic plasticity during the learning phase, that make different readouts update their connections during presentation of different subsets of patterns. However, in this paper we stick to the simple learning rule ((4.1)), and do not consider any decorrelation mechanisms. So, in the present contexts, the only way of increasing the classification capacity of the network without reaching the saturation is to increase the number of input units *N*. Also, in order to satisfy the requirement of limited connectivity, the number of connections converging onto the same readout, *C_F_* can not increase with *N*, and we need to add new readouts to connect to the newly added input units. We denote the number of readouts (number of committee members) by *M* and we derive the classification capacity *P*_max_ under the assumption that *N*, *M* and *P*_max_*f* are large numbers and the *C_F_* connections of every readout are chosen randomly and independently of any other (there will be a random overlap).

If we use the simple local learning rule (4.1), the maximum number of classifiable inputs is:

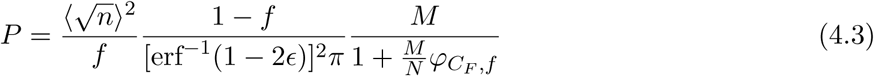

where *φC_F_,f* is of the order of *C_F_*, and depends on *C_F_* and the coding level *f*, but not on *N* or *M*. 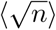 is the mean of 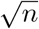 over the binomial distribution **B**(*C_F_* – 1, *f*), which is approximately 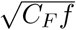 in the case *C_F_ f* ≫ 1 (dense regime), and *C_F_ f* in the case *C_F_ f* ⋘ 1 (ultra-sparse regime). Using also the approximations for *φC_F,f_* in these two cases we get (we drop the subscript _max_ from now on)

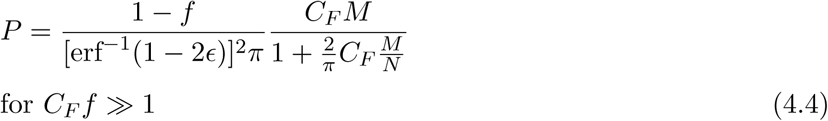

and

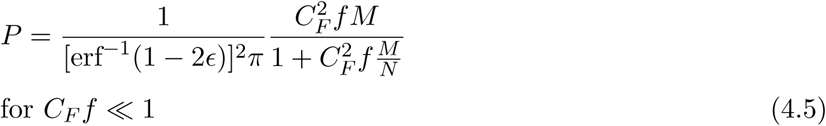

So, the dependence of *P* on the coding level *f* is weak, unless *C_F_ f* becomes smaller than 1. For sparser representations, the capacity becomes proportional to *C_F_ f*. This is not too surprising because when *C_F_ f* < 1 a significant proportion of perceptrons will read out only inactive neurons, which are not informative about the input. However, even for very sparse representations the capacity can be restored by increasing the expansion ratio *M*/*N* (see Figure 4c and 4f).

**Figure 4.**
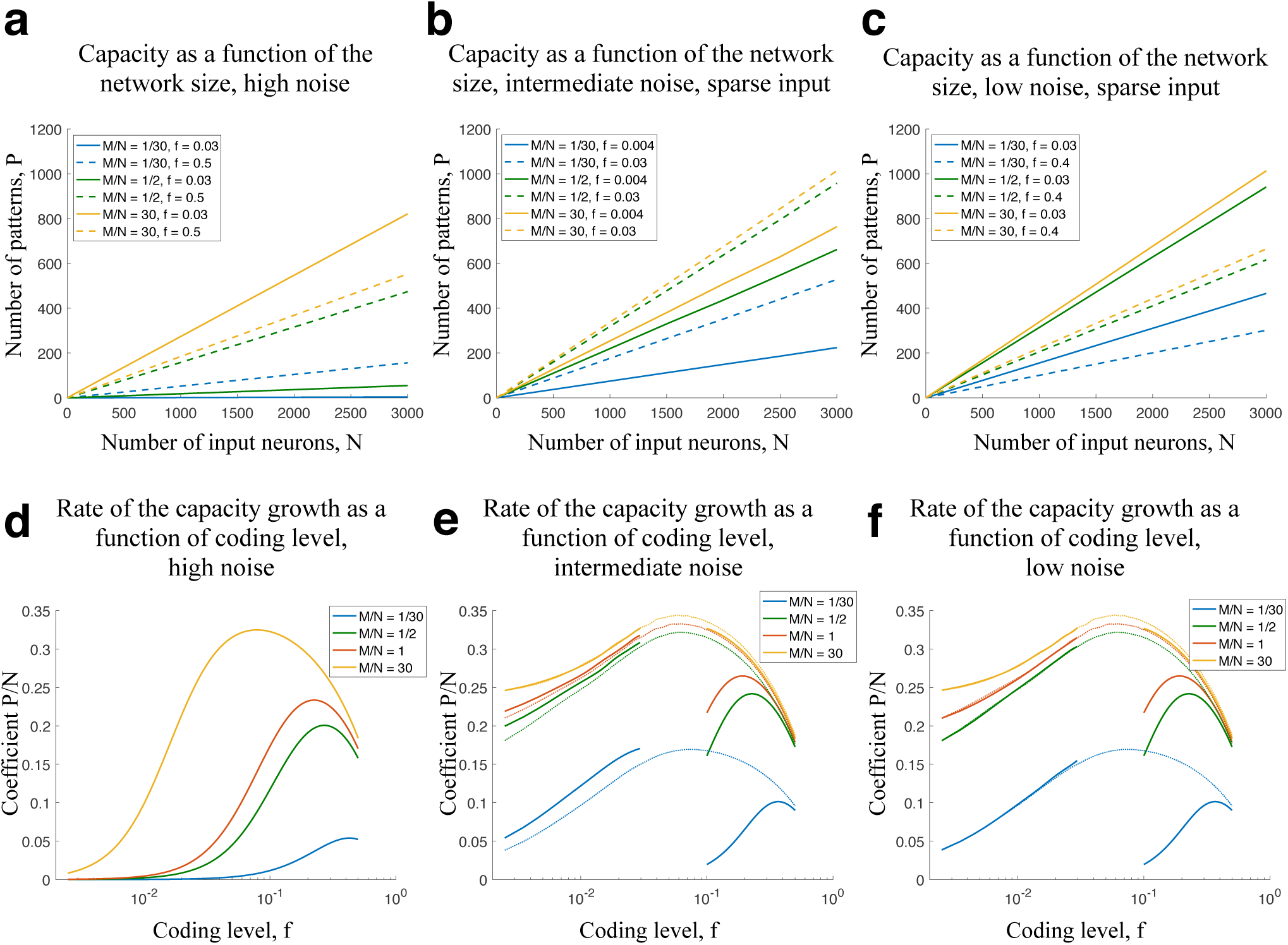
**a** – **c** The dependence of the classification capacity of the recurrent readout on the number of input units *N*, when the number of readouts *M* is increased proportionally to *N*. a High noise regime, see equation (4.6). Solid curves correspond to sparse input representation, coding level *f* = 0.03 (*C_F_ f* = 1.5) and dashed curves - to dense representation, *f* = 0.5. Different colors represent different expansion ratios *M*/*N*. The other parameters the same as in (d). **b**). Same as (a) but for the case of intermediate noise and sparse input (two-subnetworks intermediate noise regime), see formula (4.8). Solid curves: *f* = 0.004 (*C_F_ f* = 0.2), dashed curves: *f* = 0.03. The parameters are the same as for the low *f* segment of (e). **c**) Same for the low dynamical noise. Solid curves correspond to sparse input representation, *f* = 0.03, two-subnetwork low noise regime, see (4.12). Dashed curves are for dense representations *f* = 0.4 (*C_F_ f* = 20 ≫ 1) in the uniform low noise regime, see (4.7). The parameters are the same as in (f), the values of a differ for the solid and dashed curves. **d**) The dependence of the rate of the capacity growth on the coding level of input representation *f*. High noise regime, see equation (4.6). Different curves correspond to different expansion ratios *M*/*N*. The other parameters are *C_F_* = 50, *C_R_* = 700, a = 0.035, *β* = 0.05 and ε = 0.05 ε) Same for the intermediate noise regime. Different colors correspond to different expansion ratios *M*/*N*, the curves are discontinuous because different formulas are valid for sparse and dense representations. See (4.8) and (4.7). The parameters are *C_F_* = 50, *C_R_* = 700, *α* = 0.000075 for *f* < 0.03 and a = 0.006 for *f* > 0.1, *β* = 100 and ε = 0.05. The dotted curves show the committee machine result. **f**) Same as d) and e) for low noise regime. See equations (4.12) and (4.7). Parameters: *C_F_* = 50, *C_R_* = 700, α = 0.00075 for *f* < 0.03 and α = 0.006 for *f* > 0.1, *β* = 9000 and ε = 0.05. The dotted curves show the committee machine result.

When *N* and *M* grow at the same rate, the number of classifiable patterns increases linearly with N, as in the case of the fully connected single perceptron that we previously considered. However, now the connectivity of each perceptron is just *C_F_*, which does not scale with *N* or *M*. This means that it is possible to overcome the limitations of sparse connectivity. Unfortunately, this is not a satisfactory solution as it just moves the problem of limited connectivity to the readout output neuron, which now has to count the votes of all *M* perceptrons, and hence needs to be connected to *M* neurons. So again, we will need a number of connections per neuron that grows linearly with *N*. We will now propose an alternative way of implementing a commitee machine, which is based on the use of recurrent connections and it will not require a fully connected output neuron (see Figure 3).

**Figure 3.**
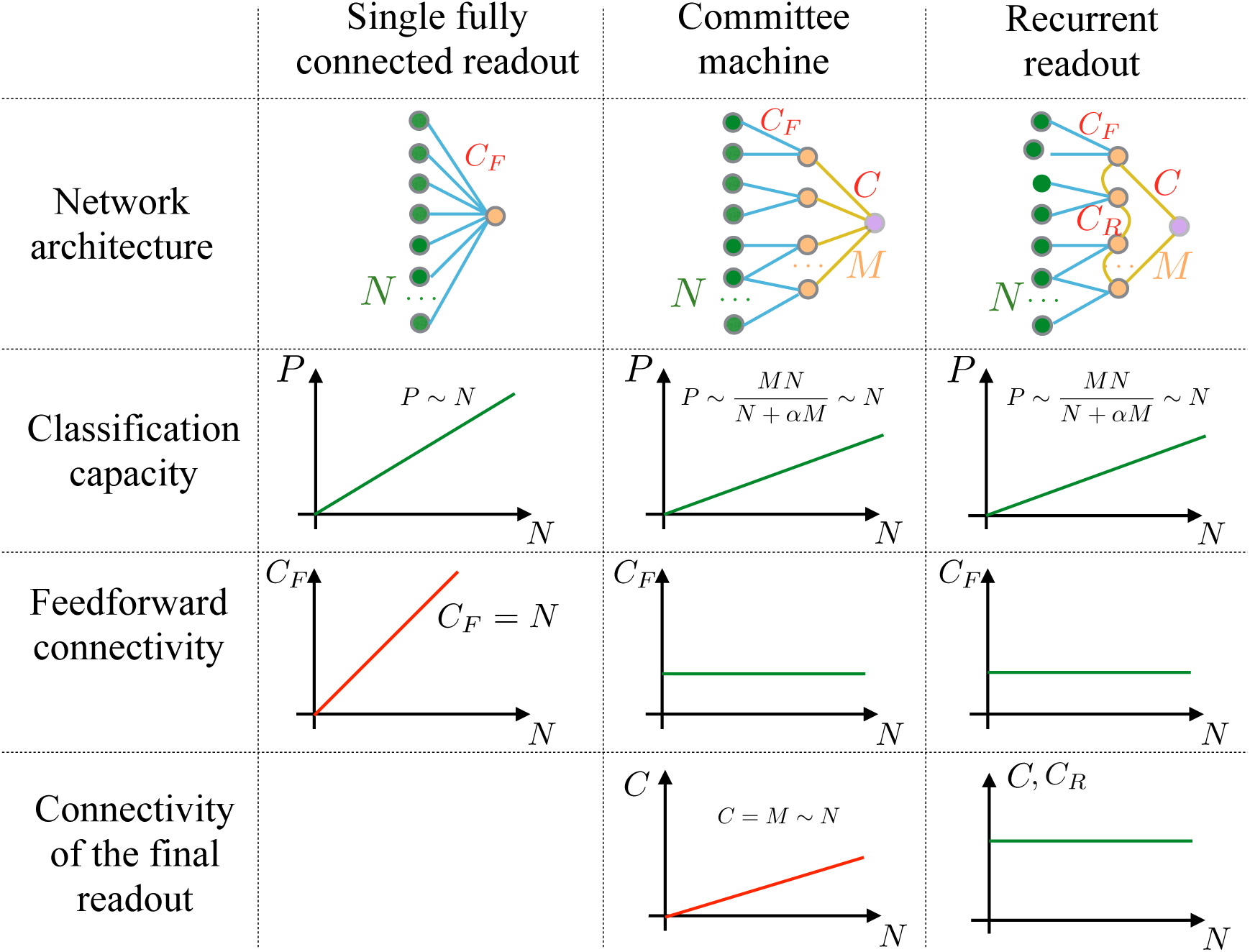
Summary of the scaling properties of the three architectures considered in the study. A single fully connected readout (classical perceptron) achieves a classification capacity *P* that grows linearly with the number of input neurons *N*, however, the number of feedforward connections that converge onto a single neuron *C_F_* also increases linearly with *N*. The Committee machine solves this problem by restricting the feedforward connectivity to a constant *C_F_*, and recovering the linear scaling of the capacity *P* by combining the decisions of *M* partially connected perceptrons via a majority vote scheme. The majority vote, however, implies a final readout, whose number of connections *C* is equal to *M*, and thus scales linearly with *N*. The aim of constructing a classifier with limited connectivity is still not met. The suggested recurrent readout architecture achieves the linear growth of the capacity while keeping *C_F_*, *C* and the number of recurrent connections per neuron *C_R_* constant as *N* increases.

### 4.2 Committee machines with recurrent connections

One way to count the votes of all perceptrons while respecting the limited connectivity constraint, would be to introduce additional layers of neurons: each neuron in the first layer would count the votes of different *C_F_* perceptrons. The neurons in the second layer would then count the votes of *C_F_* first layer neurons, and so on. For this architecture, the number of neurons would decrease by a factor *C_F_* in every new layer, leading to total number of neurons which would scale as log(*M*) or, equivalently, as log(*N*). It is also possible to set up a multi-layer network with the same number of layers in which every layer contains the same number of neurons *M*. This network would require more neurons, though it would be functionally equivalent to the first one that we considered. An interesting aspect of this architecture is that it can be interpreted as a recurrent network unfolded in time: if one assumes that the network dynamics is discrete in time, then every layer could be seen as the same recurrent network at a different time step. Importantly, the weights of the synaptic connections should be the same for every layer, as it is always the same network but at different time steps. As this network would also be functionally equivalent to the first multi-layer network that we discussed, a recurrent network can in principle replace a complex multi-layer readout which would require significantly more neurons.

These considerations induced us to study the architecture represented in Figure 1c: each perceptron of the committee machine is now connected to a randomly chosen set of the others through recurrent connections, whose weights are all the same and equal to *α*. The number of recurrent connections per perceptron is *C_R_*.

The recurrent dynamics has basically the role of stabilizing only two attractor states of the network: one in which all perceptrons are in the active state, and one in which they are all in the inactive state. These two states represent the two possible responses of the output and correspond to the two classes the input could belong to. The system is equivalent to a spin glass in the ferromagnetic state, or for a more biologically relevant analogy, to the recent decision making network of spiking neurons [33] in which only the two states corresponding to the possible decisions become stable when a sensory stimulus is presented.

Once the network has relaxed into one of the two stable states, it becomes easy to determine the class to which the input belongs, as in principle it is sufficient to read out a single perceptron. However, a single neuron readout would not be robust to noise, and hence we will consider the situation in which a number of different perceptrons are read out. We will show that this number remains finite when *N* and *M* become large, which is equivalent to saying that it is possible to construct a network, in which all the neurons, including the output neuron, have limited connectivity and the number of classifiable inputs grows linearly with *N*.

The number of classifiable inputs is derived analytically in the Methods (section 3) using a mean field approach. This number depends on the parameters that characterize the network architecture (i.e. the number and the connectivity of the different type of neurons), and on the statistics of the inputs that have to be classified. Depending on the assumptions about the parameters, there are different regimes that lead to different analytical expressions.

There are two distinct regimes that depend on whether all the recurrently connected neurons can be considered statistically equivalent or not. We call *uniform* the regime in which all the neurons can be assumed to be equivalent. This is a reasonable assumption in many situations that we discuss below, but it might not be when the number of neurons that receive no feed-forward input, which can behave differently from the others, is sufficiently large. This number is negligible when Cf *f* ≫ 1. The uniform regime is the first one that we will study systematically. Then we will discuss the non-uniform regime.

#### 4.2.1 The uniform regime

Another factor that determines the parameter regime is the amount of noise that is injected in the neurons. It is important to test the neural system in realistic conditions and to show that it is robust to noise. We introduced noise as in the Hopfield model: the state of each neuron is stochastic and its total synaptic current determines the probability distribution of the states. The noise is characterized by a parameter *β*, which in the language of statistical mechanics would be the inverse temperature parameter. When *β* is large, the noise is small and the neurons are basically deterministic. As *β* goes to zero, the neurons become more noisy and less dependent on the total synaptic input.

As we know from previous studies on attractor neural networks (see e.g. [2]), the noise cannot be too large, otherwise the attractor states remain stable only for a short time. More specifically, the noise should be smaller than the recurrent input when the network already settled in one of the two attractors and most of the presynaptic neurons are in the right state. In the uniform regime, this requirement is expressed as *β*^−1^ < *C_R_α*. Moreover, in order to guarantee attractor stability, the recurrent input should also dominate over the feed-forward one. More formally this condition can be expressed as *A* < *C_R_α*, where *A* is approximately the range in which the feed-forward synaptic input varies when different inputs are presented. It basically determines the selectivity to the inputs in the absence of the recurrent connections (see Methods for more details).

The relation between *A* and *β* is less constrained: the network architecture that we are discussing can work in different regimes that depend on how large the noise is compared to the typical amplitude of the feed-forward input.

In the *high noise* regime the noise is so large compared to the feed-forward input (*β*^−1^ ≫ *A*) that all the different recurrent neurons can behave similarly (uniform regime) even when the feed-forward input is so sparse (*C_F_ f* ≲ 1) that many neurons receive zero input. It is important to remind that in this regime the noise is large compared to *A*, but still small compared to the recurrent input. The number of classifiable patterns *P* for the high noise, always uniform regime, is given by

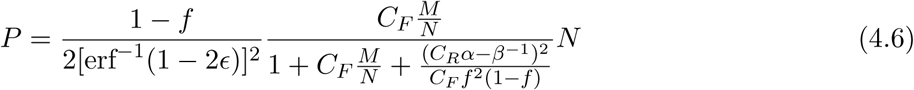

As in the committee machine case, if the number of input units *N* and the number of intermediate readouts *M* are increased in the same proportion, the number of classifiable inputs scales linearly with *M* or *N* (see Figure 4a). However, now there is not a single neuron that is required to have a connectivity that scales with *N*, the connectivity of each neuron can remain a finite number even when *N* and *M* become arbitrarily large. *α* is the strength of the recurrent connections and ε is the maximum tolerated error rate.

The rate at which *P* grows with the number of input neurons *N* (we define *P*/*N* as the capacity of the system) depends on the expansion ratio *M*/*N* (the number of intermediate readouts per input neuron), the coding level *f* and the parameters of the recurrent dynamics,*β* and *α*. The slope of the curves in Figure 4a, which represents the capacity, increases with the expansion ratio, but only up to a certain point. For *f* = 0.5 (dense representations) the capacity already saturates at *M*/*N* ≈ 1 (see the dashed lines on Figure 4a). It saturates at larger *M*/*N* for sparser representations (f = 0.03). Changing the coding level while keeping the expansion ratio fixed can either increase or decrease the capacity. The dependence of the capacity *P*/*N* on the coding level for different values of the expansion ratio is illustrated in Figure 4d.

When the noise is low compared to both the recurrent and the feedforward input, the density of the input representations starts playing a crucial role in determining whether the network is in a uniform or non-uniform regime. If the input representation is dense *C_F_ f* ≫ 1, the network is in a uniform regime. As before, all the neurons have the same average activity, but the main source of inhomogeneity is the feedforward input rather than the noise. The number of classifiable inputs in this *uniform low noise* regime is:

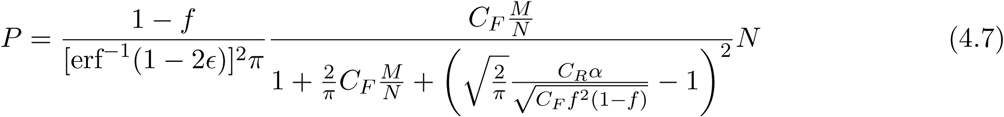

This formula is similar to one for the high noise regime. One obvious difference is that the inverse temperature parameter *β* does not appear because we assumed to be in the low noise limit *β* → ∞. The dependence of the capacity on the expansion ratio *M*/*N* is similar to one for the high noise regime. The dependence of capacity *P*/*N* on the coding level is summarized by the parts of the plots in which *f* is large in Figure 4e,f (right-hand side segments of the solid curves).

When compared to the dense limit (*C_F_ f* ≫ 1) of the majority vote result (4.4), the low noise regime formula (4.7) entails a smaller capacity. The difference comes from the last term at the denominator, which reduces the capacity. This term can be made small by tuning the parameters of the recurrent dynamics *C_R_* or *α*, but it cannot become zero, because this would correspond to the case in which the recurrent dynamics has only one stable state, which is not suitable for performing a classification task.

#### 4.2.2 Non-uniform regimes

When the noise is small compared to the feed-forward input and the representations are sparse, the uniform approximation is not valid and the recurrent network behaves in a qualitatively different way: for each input pattern, there would be two distinct populations of neurons: the *free neurons*, which receive zero feed-forward input, and hence are not constrained (free) by the input, and all the others, the *input-receivers.* The two populations would be different for different inputs, they would have different activity distributions and would evolve in time differently, although they constantly interact with each other.

Generally such a regime is intractable with the mean field method, so we need to make the additional assumption that the feed-forward synapses are sufficiently strong relative to the recurrent ones, so that, the non-zero feed-forward inputs are typically larger than the total recurrent inputs in the initial state (before the network reaches the final state when most of the neurons have the same activity). Furthermore, we need to assume that these feed-forward inputs are also much larger than the noise. Under all these assumptions, the state of the input-receivers is determined by the feed-forward input, at least in the initial stages of the dynamics, while the network is deciding which stable state to choose. We then need only to consider the dynamics of the sub-network of free units, treating the recurrent input from the input receivers as a fixed external input. It is this input, that contains the information about the correct classification.

We refer to the described scenario as to the *two-subnetworks* regime. The classification capacity in two-subnetworks scenario depends also on the noise. The noise has to be small in comparison to the feed-forward input, but it can be either small or large when compared to the amplitude of the recurrent input coming from the input-receivers. This comparison distinguishes between the *two-subnetwork low noise* and the *two-subnetwork intermediate noise* regimes.

Two-subnetwork intermediate noise regime is realized when the representations are sparse (*C_F_ f* ≲ 1) and the noise is small relative to the feed-forward input but large in the subnetwork of free neurons, namely relative to the input into free neurons from the input-receivers. This regime leads to the classification capacity of

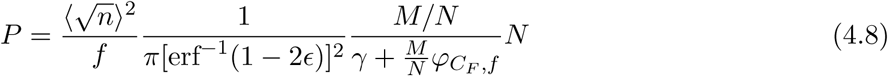

where *φC_F,f_ M*/*N* comes from the correlations between the input-receivers, *φC_F,f_* is of the order of *C_F_*, and depends on *C_F_* and on the coding level *f*. 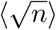 is the mean of 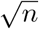 over the binomial distribution **B**(*C_F_* – 1, *f*) and *γ* is a quantity given by:

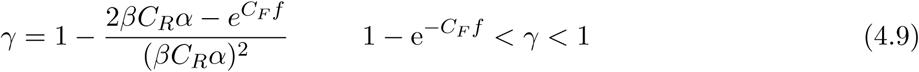

which is the smallest (highest capacity) when the network is close to transitioning from three fixed points (Figure 2c,d) to one fixed point (Figure 2a,b).

In the ultra-sparse limit, *C_F_ f* ⋘ 1 the expression for *P* becomes

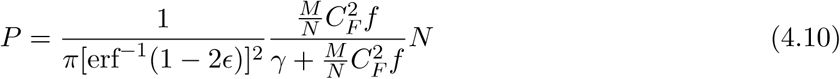

Figure 4b shows the linear dependence of *P* on the number of input neurons *N* for different expansion ratios. We can see that unless the expansion ratio is very high, even for very sparse representations (*f* = 0.004, *C_F_ f* = 0.2) the capacity grows at a similar rate or even faster compared to the case of dense representations (*f* = 0.5, *C_F_ f* = 25) in the high noise regime. The dependence of the capacity on the coding level is summarized in Figure 4e, where the curves in the low *f* region correspond to the two-subnetworks intermediate noise regime (4.8), and the segments at high values of *f* - to the uniform low noise regime (4.7). Apart from the coding level, the only parameter that differs between the two discontinuous parts of the plot for a given expansion ratio is the strength of the recurrent connections *α*. We decided to choose different as for two parts because keeping all the parameters the same while satisfying all the conditions for the two-subnetworks intermediate noise regime at low *f* and uniform low noise regime at high *f* would have required unrealistically high values of the recurrent connectivity *C_R_*. The dotted lines on the plot represent the results for committee machine with same expansion ratios.

The minimal possible value of *γ* in (4.9) is 1 – e^−*C_F_ f*^ To see this we rewrite the expression (4.9) as

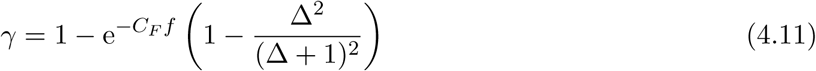

where

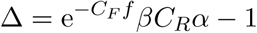

Δ must be positive in order to have three fixed points (Figure 2c,d). This implies that the expression in the paranthesis of (4.11) is positive and less than 1, from where it follows that 1 – *e^−C_F_f^* < *γ* < 1.

The lower bound for *γ* is approximately equal to *C_F_*f in the ultra-sparse limit. Plugging this value into (4.8) leads to a capacity that is basically independent from *f*. This would mean that one can decrease the coding level way below 1/*C_F_* without sacrificing the classification performance. However, keeping *γ* of the order of Cff requires having Δ of the order of 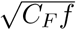 or smaller, which entails a progressively finer adjustment of the inverse temperature parameter *β* as *f* decreases. In order to have a capacity that does not become infinitesimal when *f* goes down to *f_min_*, *β* should be adjusted with the a maximum error 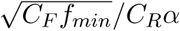.

Figure 6 (dashed lines) shows the capacity *P*/*N* as a function of *C_F_ f* for a different number of feed-forward connections per input unit *C* = *MC_F_*/*N*. Along these curves, Δ is kept at a fixed value Δ = 0.2 by choosing a new *β* for every value of *f*. Note that the curves are almost flat as long as *C_F_ f* > *C_F_f_min_* where *C_F_f_min_* ≈ Δ^2^ = 0.04

**Figure 6.**
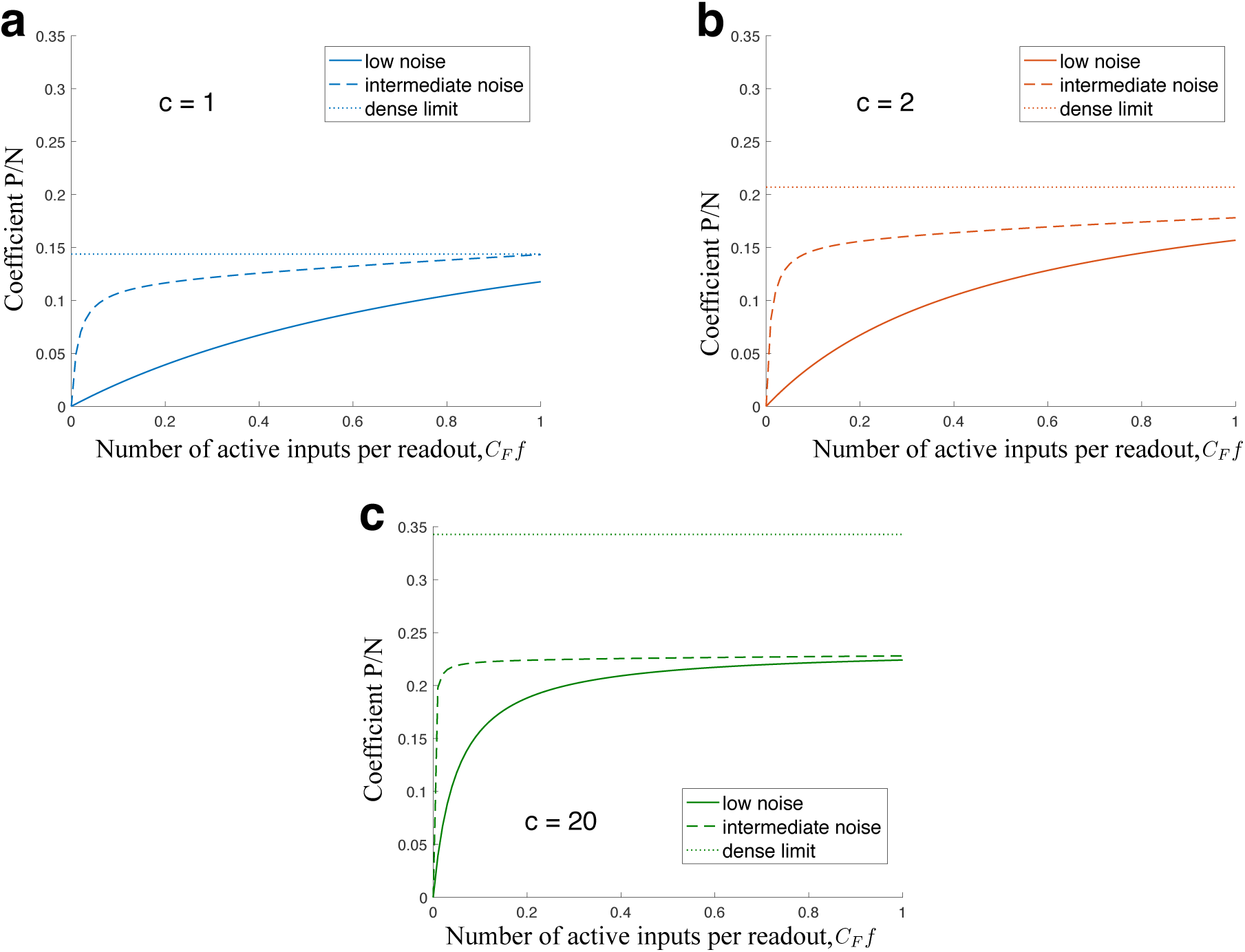
The coefficient *P*/*N* as a function of sparsity of the input representation, expressed as *C_F_ f*, for different values of the number of outgoing connections per input neuron, *c* = *C_F_ M*/*N*, in the sparse limit of two-subnetwork low and intermediate noise regimes, (4.13) and (4.10). For panel **a.** *c* =1, for panel **b.** *c* =2 and for panel **c.** *c* = 20. Solid curves correspond to the low noise regime, which is equivalent to the sparse limit of the majority vote result (see (4.13) and (4.5)), dashed lines represent the result for the intermediate amount of noise (4.10). The inverse temperature parameter *β* is different for different values of *C_F_ f* so as to keep *βαK*e*^−C_F_f^* = 1.2. The dotted lines show the maximum capacity possible for the given *c*. This is computed from the dense limit of the majority vote result (4.4), assuming that *C_F_* is large enough so that when *f* is small enough to be neglected in comparison to 1 in the numerator of (4.4), we are still in the dense regime *C_F_ f* ≫ 1.The tollerated error rate ε = 0.05.

Clearly, the capacity in the two-subnetworks intermediate noise regime is larger than in the case of a majority vote committee machine when one assume that the sparseness of the representations is the same (see (4.3) and (4.8)). This result is counterintuitive, but it can be explained: in the majority vote scenario, both the input receiving units and the free units contribute to a collective decision, even though the free units carry no information about the class of the input pattern and they actually generate noise as we assume that initially they are in a random state. In contrast, in the recurrent case, the collective state of the network is initially determined mostly by the input receiving units, which then drive the free units to the right state. The noise contained in the initial state of the free units does not affect much the initial relaxation dynamics provided that the noise in dynamics is sufficiently large (relatively low *β*).

In the case of the majority vote committee machine, the class is decided in only one time step and the initially random free units generate a certain amount of noise that depends on their number. In the case of the recurrent dynamics, the connectivity is sparse and each neuron that participates in it samples the noisy neurons a number of times that depends on the relaxation time. If these neurons can flip randomly at every time step, then their noise is averaged out and the final effect of the free units can be smaller than in the majority vote committee machine.

We now consider the two-subnetworks low noise regime. Not surprisingly, the capacity is identical to the sparse limit of the majority vote scenario (see (4.3))

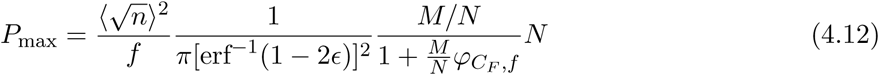

And in the sparse limit, *C_F_ f* ⋘ 1:

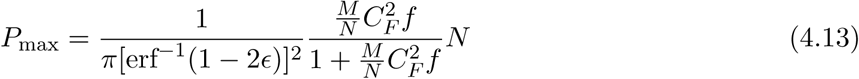

This result is summarized graphically in Figure 4c and 4f. The low *f* curve segments in Figure 4f correspond to the two-subnetwork low noise regime (identical to the result for the committee machine), and the high *f* segments - to the uniform low noise regime (4.7). The only parameter that differs between the segments of the same color is the strength of the recurrent connections *α*.

Another way to visualize the results for the two-subnetwork regime is presented in Figure 6 where we plot the capacity as a function of the product *C_F_ f*, for the two-subnetwork low noise regime in the sparse limit (4.13) (solid curves) and the same quantity for the two-subnetwork intermediate noise (4.10) (dashed curves). For the intermediate noise regime we tune the amount of noise so that *βC_R_α*e*^−C_F_f^* is always equal to 1.2. The closer this value is to 1, the larger the advantage of the intermediate noise regime over the low noise regime (majority vote). The different colors represent different values of the number of feed-forward connections per input neuron *c* = *C_F_M/N*. The dotted lines show the maximum possible capacity for a given value of c, which is achieved for the dense limit of majority vote (4.4), assuming that *C_F_* is large and *f* is small, so that *f* ⋘ 1, while *C_F_ f* ≫ 1.

It can be seen from Figure 6 that, as discussed before, the intermediate noise regime allows for very sparse input representations without sacrificing much the classification capacity, which is only slightly smaller than in the dense case (this ratio increases with *c* but saturates at *P_dense_*/ *P_sparse_* = π/2).

This means that the representations can be very sparse, despite the limited connectivity. If there is any other computational reason for preferring sparse representations, then the readout system that we propose can still be used because it can tolerate a high degree of sparseness. This might be the case of the network architecture in the hippocampus in which the representations in the dentate gyrus (DG) are extremely sparse, and the downstream readout neurons in CA3, which would be analogous to the perceptrons in our intermediate layer, have very sparse connectivity. Although we know that moderate sparseness (f ∼ 0.1) can be highly beneficial for generalization [34], we do not know why the representations in the DG are so sparse (f ∼ 0.01). However, our study shows that this elevated degree of sparseness does not necessarily impair the ability of the readout to perform efficiently a classification task.

### 4.3 Simulation Results

Figure 5 shows the results of numerical simulations run to verify the predictions of the above calcula-tions. The two plots correspond to two different regimes characterized by different coding level of the input patterns representation. Figure 5a shows the case of dense input representations *C_F_ f* = 10 ≫ 1. The theoretical predictions for the majority vote scheme (committee machine) and the recurrent readout are represented by the two dashed lines, and the solid lines with the error bars depict the results of the simulations in the two scenarios. The case of sparse input representations *C_F_ f* = 1 is shown in Figure 5b. As predicted by the analytical calculations, the recurrent readout scenario leads to higher classification capacity than the majority vote.

**Figure 5.**
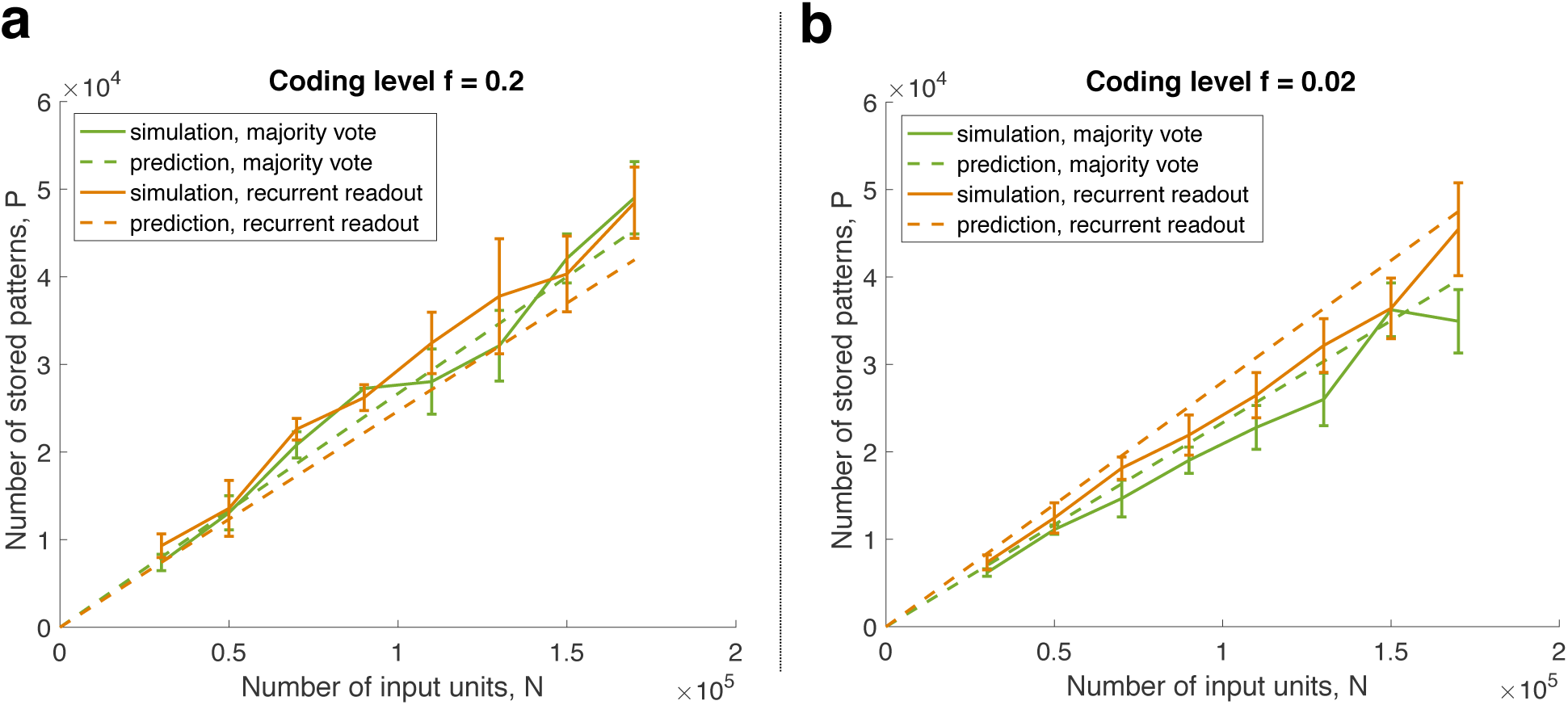
**a**. Simulation results (solid lines) and theoretical predictions (dashed lines) for the case of dense input representations, *C_F_ f* = 10. The green curves correspond to majority vote scenario (committee machine) and the orange - to the recurrent readout in the uniform regime with relatively high noise. **b**. Same for the case of sparse input representation, *C_F_ f* ≫ 1. The recurrent dynamics of the intermediate layer is in two-subnetwork regime with relatively high noise.

The simulation plots were obtained as follows: we fixed the required accuracy of the classification at 1 – ε = 0.9 and the number of feedforward connections per readout at *C_F_* = 50. For each number of the input units *N*, we chose the corresponding number of the intermediate readouts *M* = *N*/30, and we computed the predicted classification capacity *P*. We then trained the network with the set of *P*_1_ = *P* random and uncorrelated patterns, and tested the classification performance on a subset of 500 learned patterns. The obtained accuracy can be expressed as 1 – 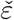. Then we trained the same network on the new set of 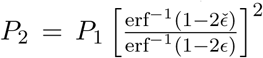 random patterns. We repeated the procedure 10 times, and we kept only those runs where the accuracy differed from the required one by no more then 2 percent (|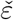 — ε| < 0.02). We then computed the mean and the standard error of the corresponding values of *P*. For the recurrent readout scenario, the recurrent connectivity was random with all the connections having the same strength, and the number of connections per unit being fixed at Cr = 200. The connectivity matrix was chosen to be symmetrical and the recurrent dynamics ran for 30 steps of synchronous update. The parameters of the recurrent dynamics for the plot of Figure 5a (dense input) were *β* = 0.5 and α = 0.015, and for the Figure 5b (sparse input): *β* = 33 and α = 0.0005

### 4.4 Optimizing the network architecture

#### 4.4.1 Optimizing the architecture under the constraint that the total number of long-range connections is constant

Now that we have derived the expression for the classification capacity as a function of the parameters of the network (4.6) - (4.13) we can determine what would be the optimal choice of parameters that maximize the networks’ performance. We first discuss the optimization under the constraint on the total number of long range connections (i.e. the feedforward connections). More specifically, we assume that the number of inputs *N* and the total number of long-range connections *C_F_M* is fixed and we ask for what value of *C_F_* (or *M*) the number of classifiable patterns *P* is maximal.

For the majority vote scenario, when *C_F_ f* is much larger than 1 (see (4.4)), *C_F_* appears in the formulae for the capacity always as a factor of *M*, so the capacity depends on the product *C_F_M*. Hence, regrouping feedforward connections, i.e. changing *C_F_* while keeping *C* = *C_F_M*/*N* constant, does not affect the classification capacity (unless we break the condition *C_F_ f* ≫ 1).

The same is true for the uniform low noise regime (4.7), which is valid only if *C_F_ f* ≫ 1. Even though *C_F_* appears in the formula in the last term of the denominator without being multiplied by *M*, it is combined with the parameters of the recurrent dynamics *C_R_α*, which can be readjusted as *C_F_* changes.

In the case of high noise and uniform regime (4.6), which is applicable for both dense and sparse representations, increasing *C_F_*, while keeping *C_F_M* constant increases the capacity. For sparser representations (lower *f*), the effect is stronger.

When *C_F_ f* ⋘ 1, and the noise is intermediate or low, we can look again at Figure 6 (see Section 4.2 and figure captions), but think of *f* as being fixed and *C_F_* as changing along the horizontal axis. In the low noise regime, the capacity decreases gradually with decreasing the connectivity *C_F_* unless the number of connections per input unit *c* is very high. Being in the intermediate noise regime, allows to go to very low values of *C_F_ f* while preserving the classification capacity.

To summarize, having less readout neurons with higher number of connections is generally better than more readout neurons with a lower number of connections. However, if the noise is not too high, this effect is not noticible unless the expected number of active inputs per readout neurons is of the order of 1 or smaller. In the intermediate noise regime, the feedforward connectivity can be decreased even further, down to 0.1 active inputs per readout, of even further if the number of feedforward connections per input neuron is high.

#### 4.4.2 Optimizing the architecture under the constraint that the total number of neurons is constant

We now determine the optimal architecture in the case when the total number of neurons is fixed. Basically we ask how to partition the total set of neurons between the input and readout layer in order to maximize the classification capacity.

It is straightforward to derive the optimal expansion ratio *M*/*N* from the formulas (4.6) - (4.13) under the constraint *M* + *N* = *const*. In the uniform regime (see equations (4.6) and (4.7)), if the parameters of the recurrent dynamics are not too far from the optimal ones, the expansion ratio that maximizes the capacity can be approximated by

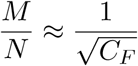

This corresponds to more than one perceptron for every *C_F_* input in the non-overlapping design (i.e. when each input neuron is connected to only one perceptron and hence the populations of input neurons that are readout by each unit are not overlapping). To be more precise, a typical input is connected to approximately *C_F_M*/*N* œ 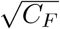 perceptrons. However, the number of perceptrons *M* is still much smaller than the number of inputs *N*.

For the two-subnetworks low noise regime in the sparse limit *C_F_ f* → 1, the optimal expansion ratio is approximately

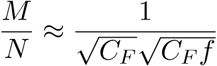

This corresponds to 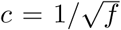 connections per input unit. For sparser representataions the optimal proportion of units in the readout layer increases.

For intermediate noise

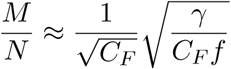

where *γ* is given by (4.9)

So, when the metabolic cost of incorporating new units into the network is high, and the number of feedforward connections converging onto the same readout unit (*C_F_*) is fixed (for example, because of spatial constraints), there is an optimal ratio between the number of input and readout units. In the case of dense input representations or high dynamical noise, this ratio is determined by the number of converging connections, *C_F_*, whereas for sparse representations - by the coding level *f*. In both cases the number of outgoing connections per input neuron is large. For dense representations the optimal architecture is convergent (*M* < *N*), while for sparse representations it can be either convergent or divergent.

### 4.5 Multinomial Classification

We now turn to a more difficult problem of classifying the inputs into more than two categories. The scheme presented above can be generalized in a straightforward way to serve as a multinomial classifier. We first show that in the case of multiple classes the classification capacity of the network does not change substantially. We later discuss a more realistic scenario of multiclass classification for which we can not compute the capacity analytically, but we demonstrate with simulations that the capacity decrease is moderate and, most importantly, that the linear scaling with the network size is preserved.

#### 4.5.1 Structured output

The immediate generalization of the recurrent readout scheme to multinomial classification task is to introduce several populations of intermediate readout neurons, each of which would correspond to one class. The recurrent connectivity within a population would be as described before, while no recurrent connections would exist between the neurons belonging to distinct populations. The desired output pattern in response to an input from each class is then structured so that the population corresponding to the given class is active while the others are inactive. The final readout has to contain multiple readout units, one for each class. Their connectivity can still be sparse and random, but the sign of the connections would have to be adjusted based on whether it comes from the neuron in the population selective for the same class as the given final readout or not (see Figure 7a).

**Figure 7.**
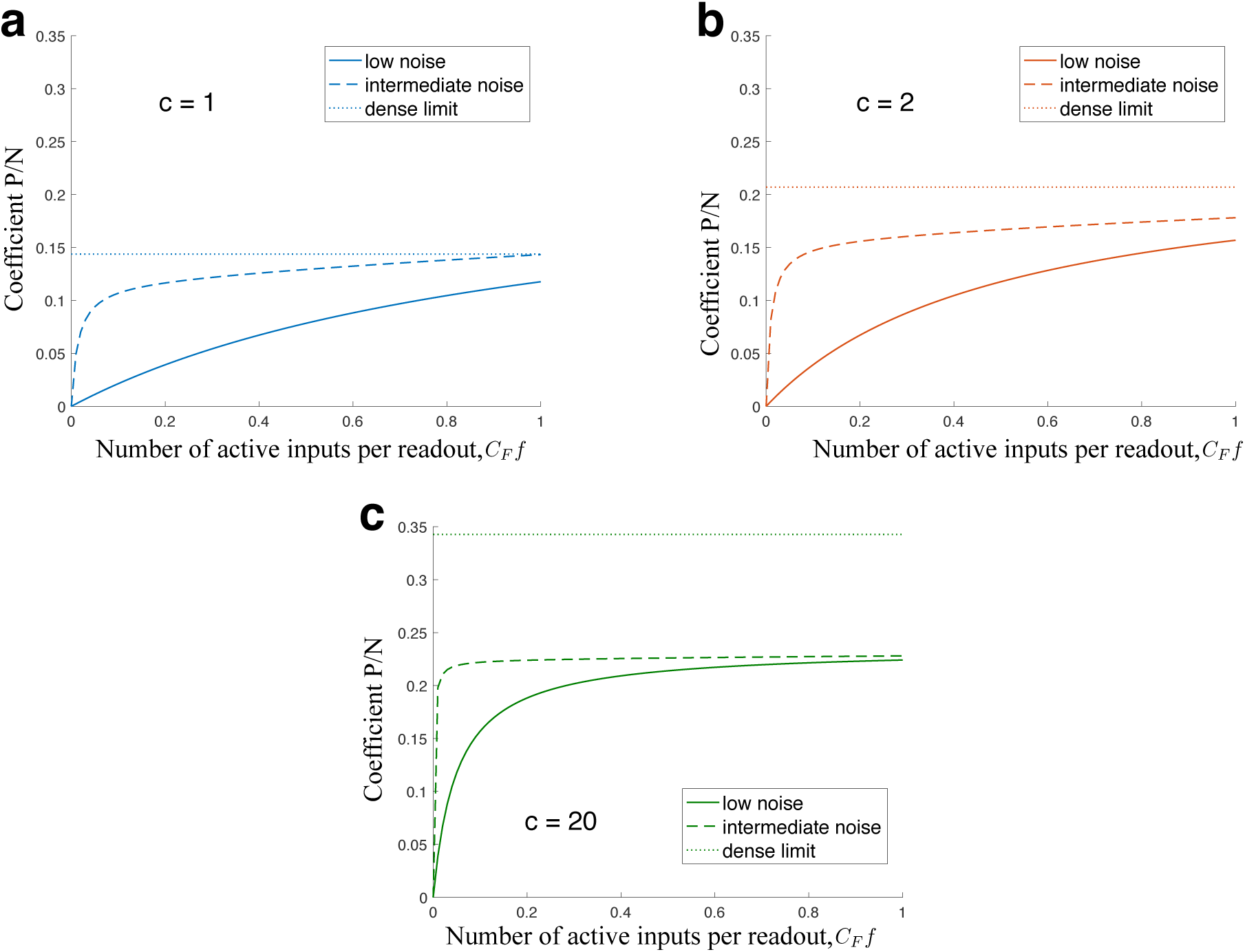
**a.** Network architecture for the case of structured output (see Section 4.5.1). For the case of 3-way classification, the intermediate layer of readout neurons is divided into 3 subpopulations, each selective for its own class of input patterns. The recurrent connectivity is random and excitatory within subpopulations, but there is no recurrent connections between the subpopulations. The final readouts, one for each class, are connected sparsely and randomly, as before, but the sign of the connections is only positive if the presynaptic neuron belongs to the correct subpopulation, the rest are zero or negative. **b.** Network architecture for the case of random output (Section 4.5.2). There are no distinct subpopulations in the intermediate layer, and the desired output pattern corresponding to each class of input patterns is chosen randomly. The recurrent connections exist between any pair of the readout neurons with equal probability. The strength of these connections, however, is now adjusted according to Hebbian learning rule (4.14). **c.** The results of the simulation for multinomial classification. The output patterns corresponding to *L* = 5 classes are chosen randomly with the coding level *y* = 1/2. The recurrent connectivity is sparse and the strength of the synapses are trained with the learning rule (4.14). The network of recurrently connected perceptrons is in the high noise regime with dense input representations (*C_F_* = 50, *f* = 0.2, *C_R_* = 200, *α* = 0.015, *β* = 0.5). The dashed line is the estimation of the capacity from the formula (4.6) assuming two equal size subpopulations of the readouts.

The classification capacity can now be computed in the same way as above, by noticing that each population is now doing a binary classification, selecting for one out of *l* classes. The only difference is that the proportion of ‘positive’ patterns (the output sparseness) is now *y* = 1/*L* instead of 1/2. The capacity formula for the case of sparse output is derived in the Methods (section 3) and it differs from the capacity for a dense case by a factor, that depends on *y*.

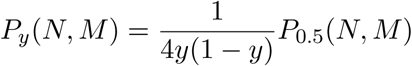

It should be noted, that the number of intermediate readouts *M*, entering this formula is the number of units in the population selective for a particular class. So, if total number of intermediate readout units is *Mtotai,* and all populations have equal size, it is *M* = *M_total_/L* = *yM_total_*, that should enter the formulas for the capacity. So, in terms of the total number of intermediate units, in the two-subnetworks intermediate noise regime, for example (formula (4.8)), we have

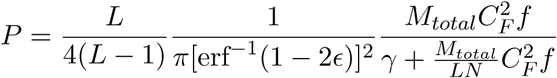

Where *γ* is given by (4.9). There are two differences with respect to the binary classification case (4.8). The first is the prefactor, which is equal to 1/2 for the case of two classes (*L* = 2). This is the reflection of the fact, that when only two classes are possible, the current scheme is redundant - when the first population is active, the other is not, and vice versa. In the limit of large number of classes, the prefactor is equal to 1/4. The other difference is in the second term in the denominator, which rescales *N*, the number of the input units. Namely, for the number of intermediate readout units, the role of correlations between them is decreased compared to the binary classification case. This is because there is no interference between the readout neurons belonging to different populations.

#### 4.5.2 Random output

Another, more realistic scenario is to assign the output patterns that correspond to each of *L* classes randomly and train the existing recurrent connections with a plausible learning rule.

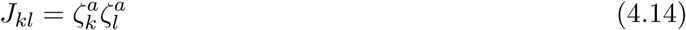

Where the 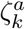 is the output pattern corresponding to class *a*, (*a* = 1… *L*). In this case there are no structurally distinct subpopulations of the intermediate readout neurons which are defined a priori (see Figure 7b). In contrast, the subpopulations of neurons which represent different classes emerge as a consequence of the learning rule (4.14).

Figure 7 shows the simulation results for a 5-way classification (*L* = 5) of dense input patterns with high dynamical noise (the parameter values are given in the figure caption). The dashed line on the figure indicates the capacity given by the formula (4.6) assuming that the population of *M* intermediate readouts is split into two segregated subpopulations, whose activity is opposite in all the output patterns.

### 4.6 The initial condition of the recurrent network

An important assumption that we made in order to implement the majority vote with a recurrent readout is that the recurrent network initial condition is unbiased, or, in other words, that *m*_0_ = 0 (see Section 3.3.3). This condition might sound difficult to realize in a network that is basically designed to amplify any small deviation from *m*_0_ = 0. However, this condition could be realized as follows: assume that before a pattern to be classified is presented, the input layer is spontaneously active. This spontaneous activity generates a feed-forward input 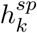 which causes the disordered state (*m* = 0) to be the only stable state of the recurrent network (see Figure 2a). There are two conditions on the statistics of 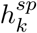 that are required to have *M* = 0 as the only stable state of the system in the mean field approximation. The first requirement is that 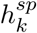 has zero expectation value, which is satisfied if the patterns of spontaneous activity are not correlated with the training patterns. The second requirement is that the standard deviation of the distribution is large enough, to make the slope of the sigmoidal curve of Figure 2 smaller than 1. For instance, in the uniform regime (see section 3.3.5), the latter requirement is 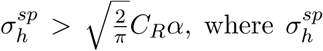 is the standard deviation of the feedforward current due to spontaneous activity. When the input pattern is presented then the noise is assumed to decrease to restore the conditions (3.47) that allow the recurrent network to have three solutions, two stable, corresponding to the possible classification outcomes, and one unstable, which was the initial state. A reduction in noise during stimulus presentation has been observed in [35].

## 5 Discussion

We presented a model network based on perceptrons in which all the neurons have limited connectivity and nevertheless the classification capacity grows unboundedly and linearly with the size of the network.

The limitations on classification capacity of the individual perceptrons that are imposed by the limited connectivity are overcome by reading out multiple perceptrons, as in a committee machine. However, the readout mechanism is different from the one normally used in committee machines as it uses a recurrent attractor dynamics of committee members to generate a final vote. Thanks to the recurrent dynamics, it is then possible to readout a small sample of all the committee members to determine the committee decision. This allows for readouts that have a limited connectivity, even when the size of the network becomes very large.

Interestingly, there are situations in which the proposed recurrent readout scheme can outperform classical readouts that are based on a majority vote despite the fact that the majority vote would require a significantly larger readout connectivity (see sections 4.2.2 and 3.3.9). For the majority vote scheme, the classification capacity drops drastically when the input representations are very sparse because the fraction of classifiers whose inputs are all silent becomes substantial and these classifiers just contribute to the noise. Instead, for the recurrent readout the classification capacity can be kept high even for very sparse representations in certain parameter regimes because the recurrent dynamics can align the ‘free’ classifiers to the majority decided by the other, informative classifiers. The lower limit on the coding level *f*, below which the capacity drops is determined by the amount of noise in the recurrent dynamics, the expansion ratio and the number of feed-forward connections per perceptron (see Figure 4).

In general, the proposed system is robust to both sparse connectivity and sparse representations, which makes it suitable to describe neural circuits like the dentate gyrus (DG) and CA3 area, where the number of connections of downstream neurons (CA3) is much smaller than the number of neurons in the input (DG) and the neural activity in the input can be very sparse. CA3 is known to have the recurrent connections that would implement our proposed readout mechanism. We showed that for intermediate noise (see sections 4.2.2 and 3.3.9), the classification capacity stays within a reasonable range even when the expected number of active units read out by each perceptron is smaller than 1 (see Figures 4e, 4f and 6). This result nicely complements the study presented in [34], where the authors show that low dimensional correlated inputs require an intermediate layer of neurons (randomly connected in [34]). For these neurons in the intermediate layer there is an optimal sparseness level which minimizes the generalization error of a single perceptron-like readout. Here we showed that there is a readout scheme that would work also for the sparse representations required in the intermediate layer of [34].

## 6 Appendix

### 6.1 A1

In this section we derive (3.43), the variance of the feedforward current, averaged over the readouts

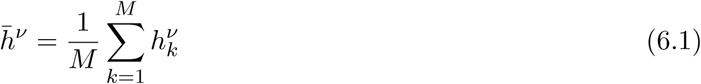

The variance of *h̅^ν^* is contributed by the diagonal terms and the non-diagonal terms, in the limit *M* → ∞ approximated by

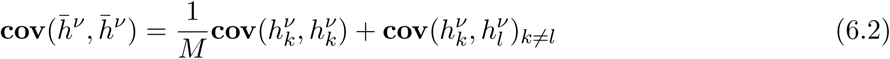

For the non-diagonal terms, neglecting 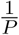 corrections coming from the signal, using representation (3.21), we find in the same way as in the computation of (3.22)

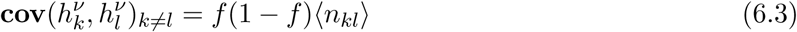

from the covariance of the *z_kl_* terms. Here 〈*n_kl_*〉 is the expectation value of the number of common active input neurons for readouts *k* and *l* (see details in section 3.2.2). Therefore, comparing with (3.9) we find

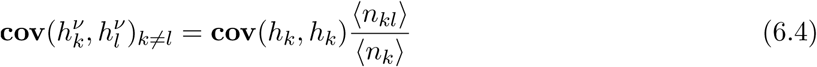

The 〈*n_kl_*〉 in the limit *N*, *M* → ∞ to and finite *C_F_*, *f* can be estimated as

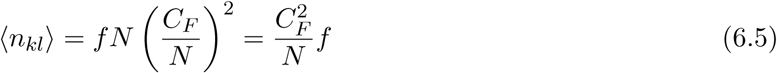

which gives

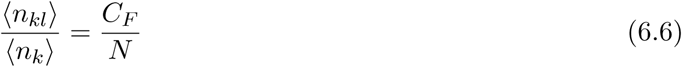

and therefore

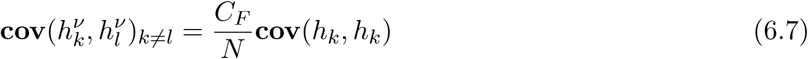

Hence (6.2) reduces to

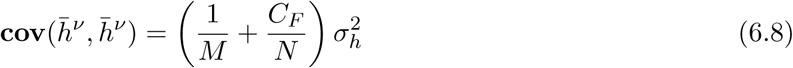

and implies (??).

### 6.2 A2

In this section we derive the formula (3.75) for the covariance of the signs of the external currents into two different free units in the two-subnetwork regime.

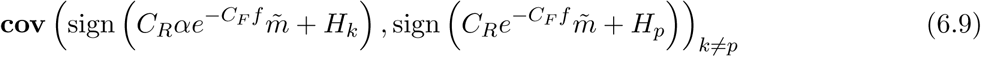

we introduce a notation

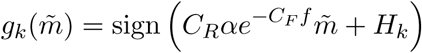

and without loss of generality assume *α* =1.

There are two cases of contributions to the correlation, that we will call case I and case II, see figure 8.

**Figure 8.**
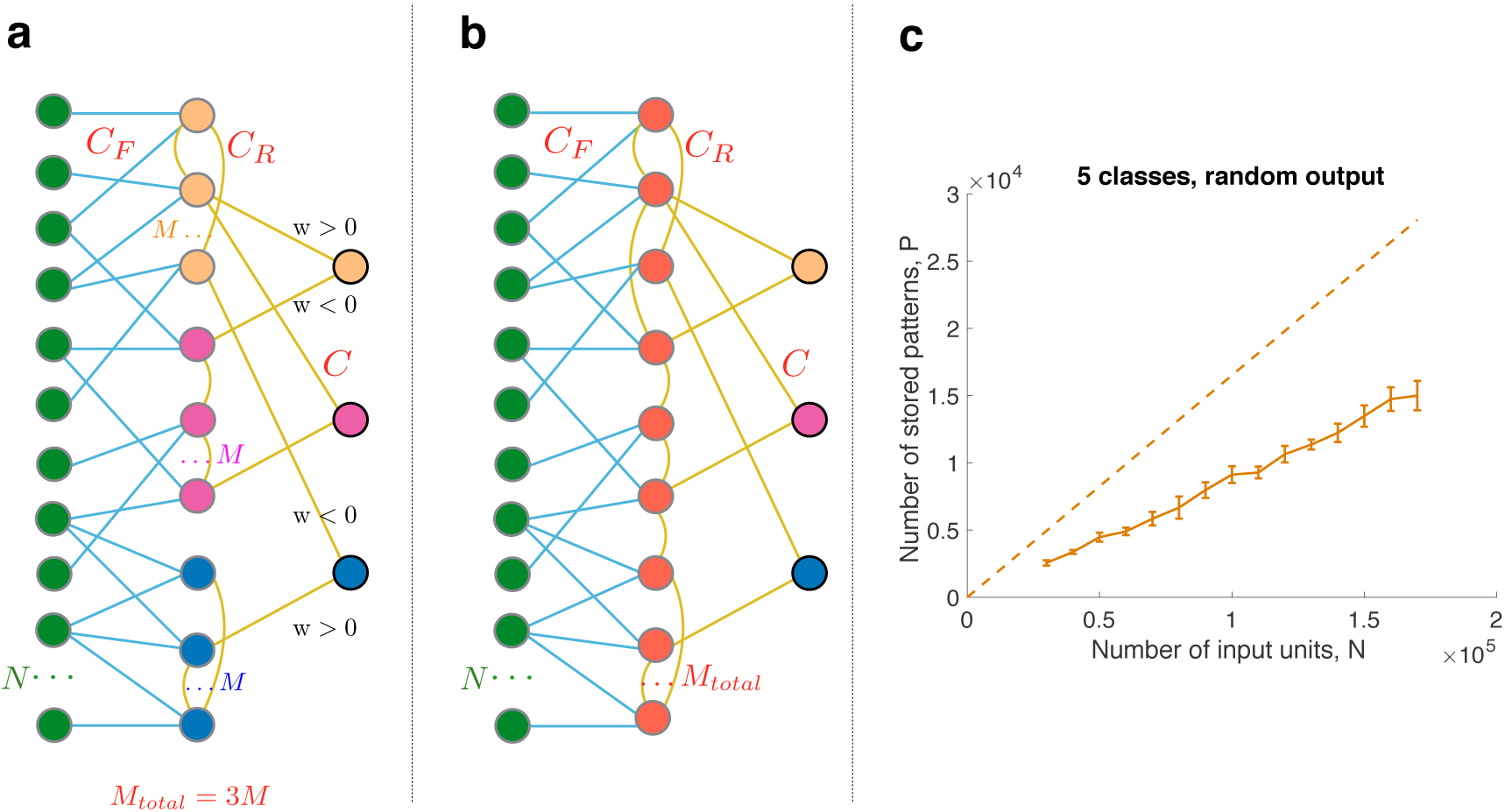
Two sources of input correlations for the subnetwork of free units (orange circles), referred in the text as case I and case II. On the left diagram two free units are connected to the same input receiving unit in the readout layer (red circle). On the right diagram there is no input receiving unit that is connected to both free units, but the correlation arises from an active unit in the input layer (green circle), which is connected to the two free units indirectly.

The case I contribution to this correlation comes from the free units *k* and *p* being connected to the same input receiving unit *r*. We neglect the probability that the overlap will be over more than one input receiving unit since we keep connectivity *C_R_* fixed when we scale the number of units *M*. To the leading order in *C_R_*/*M*, the probability of case I contribution is

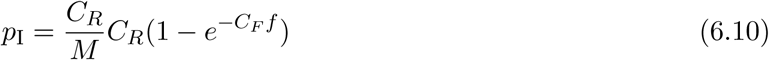

This is because a typical free unit is connected to *C*_R_(1 – *e^−C_F_f^*) out of *M*(1 – *e^−C_F_f^*) input receiving units.

The case II contribution comes from the possibility that there is an input layer unit that is active and connects to both via different input receiving neurons. The approximate robability of case II contribution, assuming *C_F_/N* is small, is given by

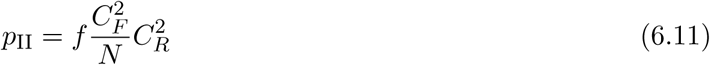

To derive this probability, recall that the probability of any two readouts to be connected to the same active input is 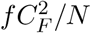, and there are 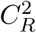 pairs of readouts (red units on figure 8) connected to the given pair of free units (orange units). The probability that both units in this pair are input receiving units is already taken into account by the factor 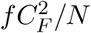.

In the case I the relevant correlation is

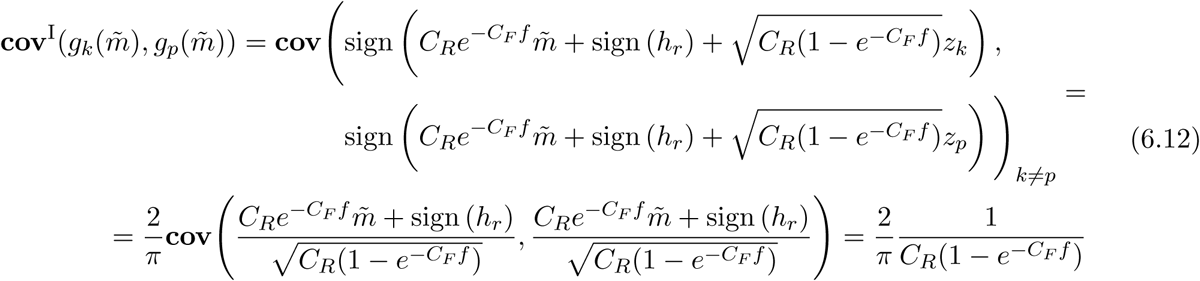

We approximated *C_R_* – 1 by *C_R_* and used error function integral (3.13) at small argument on standard Gaussian variables *z_k_* and *z_p_* to transform the first line to the second line.

In the case II the relevant correlation

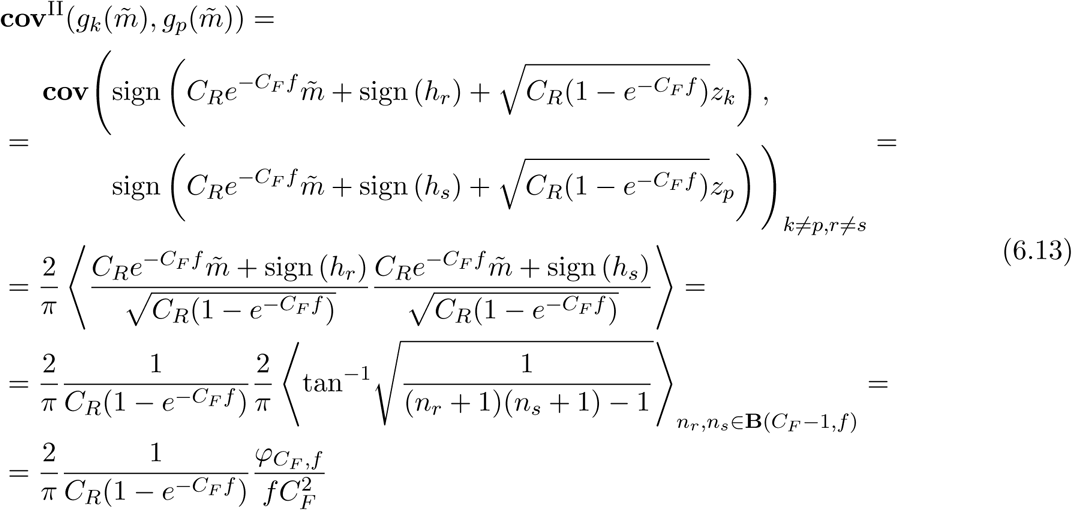

where *n_r_* and *n_s_* are from binomial distribution on *C_F_* – 1 trials with probability *f* computed as in (3.25) from the correlation of the sign(*h_r_*) and sign(*h_s_*).

Now we can compute 6.9 in the leading order as *p_I_, p_II_* probability weighted sum of the contributions from case I and case II:

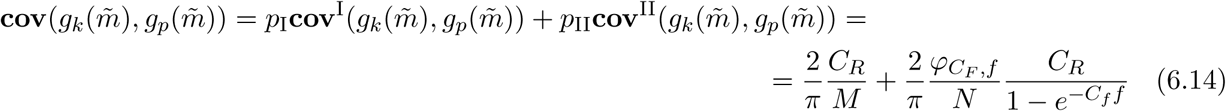

At the diagonal terms we have simply

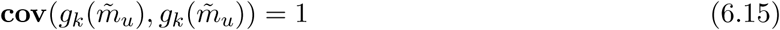

Alltogether, combining the contribution from diagonal and non-diagonal terms as in (6.2) we find

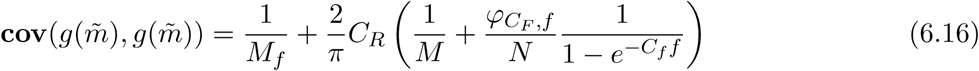

We will assume that *C_F_ f* ≲ 1 so that Mf ≃ *M* and that even though *C_R_* does not scale linearly with *M*, *N*, *P* still

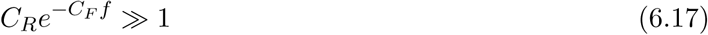

then we can, in fact, drop the diagonal term in (6.16) and take the approximation

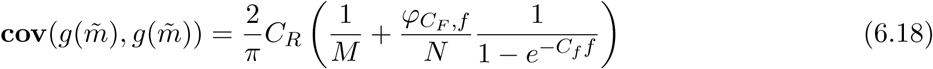

## 7 Acknowledgements

SF is supported by the Gatsby Charitable Foundation, the Simons Foundation, the Schwartz foundation, the Kavli foundation and the NSF’s NeuroNex program award DBI-1707398. LK acknowledges support from the grants ANR-10-LABX-0087 IEC, ANR-10-IDEX-0001-02 PSL.

## Footnotes

1 See equation 18 on page 158 in [29]

